# Genomic instability is an early event driving chromatin reorganization and escape from oncogene-induced senescence

**DOI:** 10.1101/2020.12.20.423639

**Authors:** C Zampetidis, P Galanos, A Angelopoulou, Y Zhu, T Karamitros, A Polyzou, I Mourkioti, N Lagopati, R Mirzazadeh, A Polyzos, S Garnerone, EG Gusmao, K Sofiadis, DE Pefani, M Demaria, N Crosetto, A Maya-Mendoza, K Evangelou, J Bartek, A Papantonis, VG Gorgoulis

## Abstract

Oncogene-induced senescence (OIS) is an inherent and important tumor suppressor mechanism. However, if not timely removed via immune surveillance, senescent cells will also present a detrimental side. Although this has mostly been attributed to the senescence-associated-secretory-phenotype (SASP) of these cells, we recently proposed that “escape” from the senescent state represents another unfavorable outcome. Here, we exploit genomic and functional data from a prototypical human epithelial cell model carrying an inducible *CDC6* oncogene to identify an early-acquired recurrent chromosomal inversion, which harbors a locus encoding the circadian transcription factor BHLHE40. This inversion alone suffices for BHLHE40 activation upon *CDC6* induction and for driving cell cycle re-entry and malignant transformation. In summary, we now provide strong evidence in support of genomic instability underlying “escape” from oncogene-induced senescence.

**HIGHLIGHTS:**

- Oncogene driven error-prone repair produces early genetic lesions allowing escape from senescence
- Cells escaping oncogene-induced senescence display mutational signatures observed in cancer patients
- A single recurrent inversion harboring a circadian TF gene suffices for bypassing oncogene-induced senescence
- Chromatin loop and compartment remodeling support the “escape” transcriptional program

## INTRODUCTION

According to the DNA damage model for cancer development, activated oncogenes trigger genomic instability that at some point breaches the tumor-suppressing barriers of apoptosis and senescence to promote cancer development [Halazonetis et al., 2008]. This model readily explains how emerging genomic instability in cancer leads, via the accumulation of inactivating mutations at key signaling hubs and regulatory factors, to evasion of apoptosis and “bypass” of senescence [Halazonetis et al., 2008; Negrini et al., 2010; Gorgoulis et al., 2018]. It also provides the grounds for considering senescence as an inherent barrier to tumour development in precancerous stages [Bartkova et al., 2006; Di Micco et al., 2006; Collado et al., 2005; Braig et al., 2005; Michaloglou et al., 2005; Chen et al., 2005]. However, this model does not explain how cells that have entered such a state of irreversible cell cycle arrest become able to breach this barrier and re-initiate proliferation.

Recently, we and others demonstrated how, under certain conditions, a subset of cells in a senescent population do re-enter the cell cycle, thus “escaping” senescence. Such “escapee” cells adopt a more aggressive phenotype that closely mimics cancer development [Galanos et al., 2016; Patel et al., 2016; Milanovic et al., 2018; Komseli et al., 2018; Yu et al., 2018; Gorgoulis et al., 2019]. Nevertheless, the molecular mechanism underlying this “escape” phenomenon has not been deciphered.

Here we hypothesize that, if our cancer development model [Halazonetis et al., 2008] should also pertain to the “escape” phenomenon, then accumulating DNA damage traits during oncogene-induced senescence (OIS) will be selected and should appear in “escape” cells as functionally meaningful genetic defects. To address this, we combine a prototypical OIS cellular system with genomics and functional assays to present the first evidence in support of this hypothesis, while also discussing its clinical significance.

## RESULTS

### An oncogene-induced senescence model recapitulating cancer evolution

We recently described a cellular system based on normal human bronchial epithelial cells (HBECs) carrying a *CDC6*-TetON overexpression cassette [Moreno et al., 2016; Komseli et al., 2018]. HBECs are of epithelial origin like most common cancer types, and in their un-induced state (“OFF” in **Figure 1A**), they are free from the mutation burden found in cancer cells [Goodspeed et al., 2016; Stratton et al., 2009]. This permits accurate detection of amassing DNA alterations during *CDC6*-induced senescence (“ON” state in **Figure 1A**).

**Figure 1.**
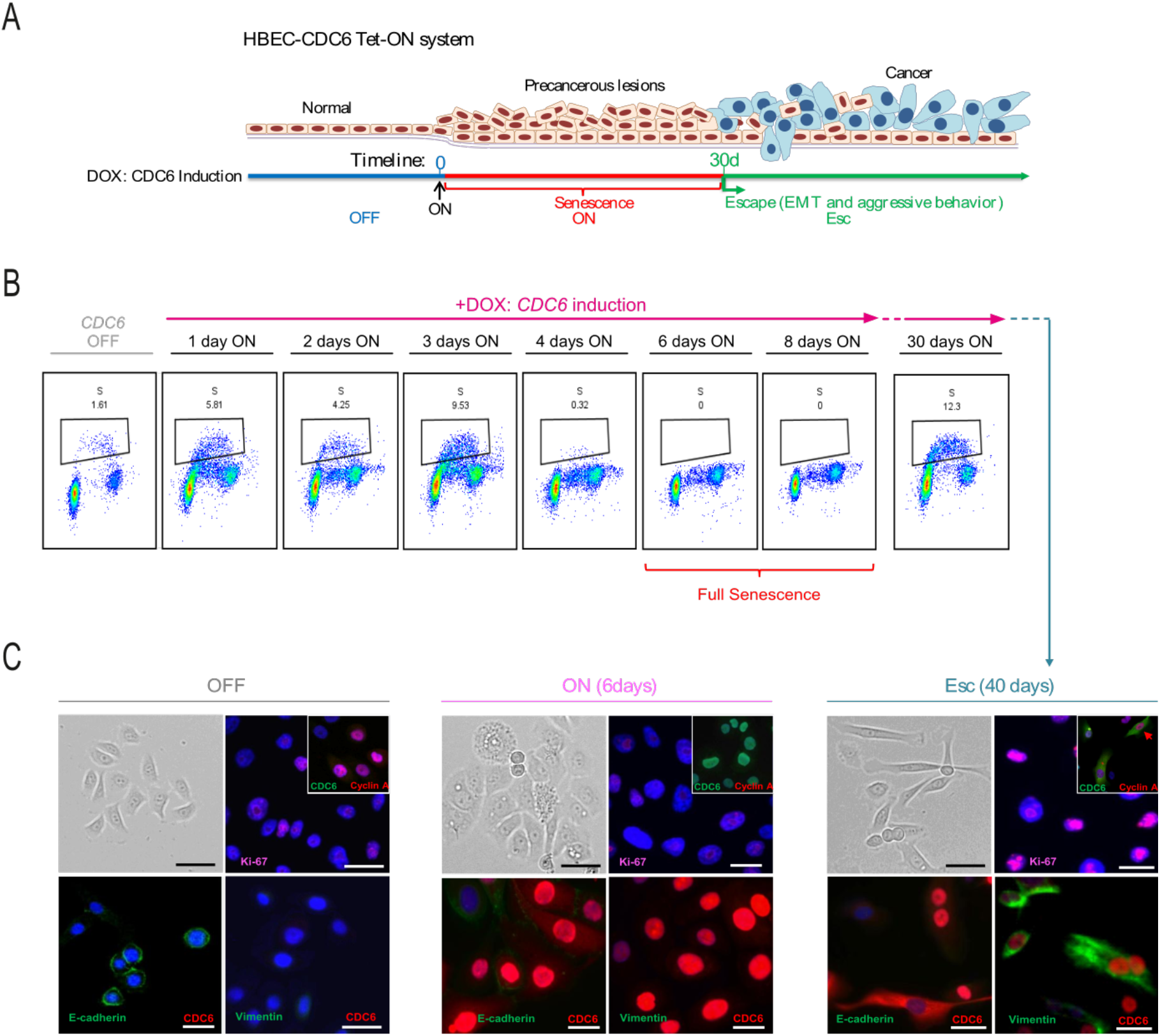
Escape from oncogene-induced senescence. (**A**) A human bronchial epithelial cell (HBEC) *CDC6*-TetON cellular system recapitulating successive stages of cancer evolution [Komseli et al., 2018]. (**B**) FACS-based cell cycle analysis of HBECs at different time points following *CDC6* induction demonstrating progressive S-phase reduction until day 4. Bar graphs show mean reduction (±S.D.; n=3). On day 25 a small number of S-phase cells reappears. (**C**) Representative phase contrast views and immunodetection of epithelial (E-cadherin) and mesenchymal markers (vimentin) in HBECs showing that escape from CDC6-induced senescence (ESC) coincides with epithelial-to-mesenchymal transition.

The replication licensing factor CDC6 was chosen as the inducible oncogenic stimulus because (i) as a key component of the replication licensing machinery that integrates most mitogenic and oncogenic stimuli, it is frequently deregulated already in the earliest stages of cancer [Karakaidos et al., 2004; Liontos et al., 2007; Sideridou et al., 2011; Petrakis et al., 2016]; (ii) compared to other oncogenes tested, such as *RAS* or *BRAF*, it proved a more powerful inducer of senescence [Patel et al., 2016]; and (iii) its overexpression is linked to poor patient survival across common cancer types (**Figure S1**).

Importantly, this system offers the advantage of prompt and quantitative senescence entry (within <6 days), followed by escape from senescence in a relatively short time period (within ∼40 days; “ESC” in **Figure 1A**) [Moreno et al., 2016; Komseli et al., 2018]. These transitions recapitulate the whole evolution course of malignant transformation, and are equally observed in 2D and 3D organotypic cell culture conditions (**Figure S2A-C**). Thus, for our working hypothesis (see **Introduction**) to be validated, the following need to occur. First, shutting off *CDC6* overexpression in cells that have “escaped” senescence should not result in phenotype reversal, suggesting acquisition of permanent molecular alterations. Second, following *CDC6* induction, DNA double strand breaks (DSBs) should form and, at least a fraction of them, should be repaired in an error-prone manner. Finally, genomic alterations produced in the senescent state should be selected to functionally facilitate “escape”.

### CDC6 expression is dispensable after EMT-like “escape” from senescence

To exclude mapping of stochastic alterations, we conducted three independent evolution experiments (**Figure S3A**). In all three, a fraction of cells (∼50 colonies/3×10^6^ cells) re-entered the cell cycle after a protracted CDC6-induced senescent phase (**Figure 1B** and **Video S1A-B**). These “escape” (ESC) cells adopted epithelial-to-mesenchymal transition (EMT) features (**Figures 1C, S2A-C** and **Video S1A-B**) known to facilitate cancer invasion and metastasis [Nieto et al., 2016; Thiery et al., 2009]. Moreover, conducting a bioinformatic analysis we observed that “escape” cells exhibited a mixed stem cell signature encompassing embryonic, epithelial, mesenchymal-like and MYC-dependent markers [Ritschka et al., 2017; Wong et al., 2008; Kim et al., 2010; Ivanova et al., 2002; Chambers et al., 2007; Milanovic et al., 2018] (**Figure S2D**). Critically, switching off *CDC6* overexpression does not result in ESC phenotype reversal, hence the preservation of growth and invasion capacities (**Figure S2E-G**).

### DSBs occur early upon senescence entry and are repaired in an error-prone manner

To determine whether and to which extent double strand breaks (DSBs) occur, we performed BLISS (Breaks Labeling *In-Situ* and Sequencing) [Yan et al., 2017] at different times after *CDC6* overexpression. BLISS analysis verified DSB emergence in senescence, with a dramatic increase after 3 days and an almost 50% reduction at the peak of senescence (day 6) suggesting that a repair process took place (**Figure 2A**).

**Figure 2.**
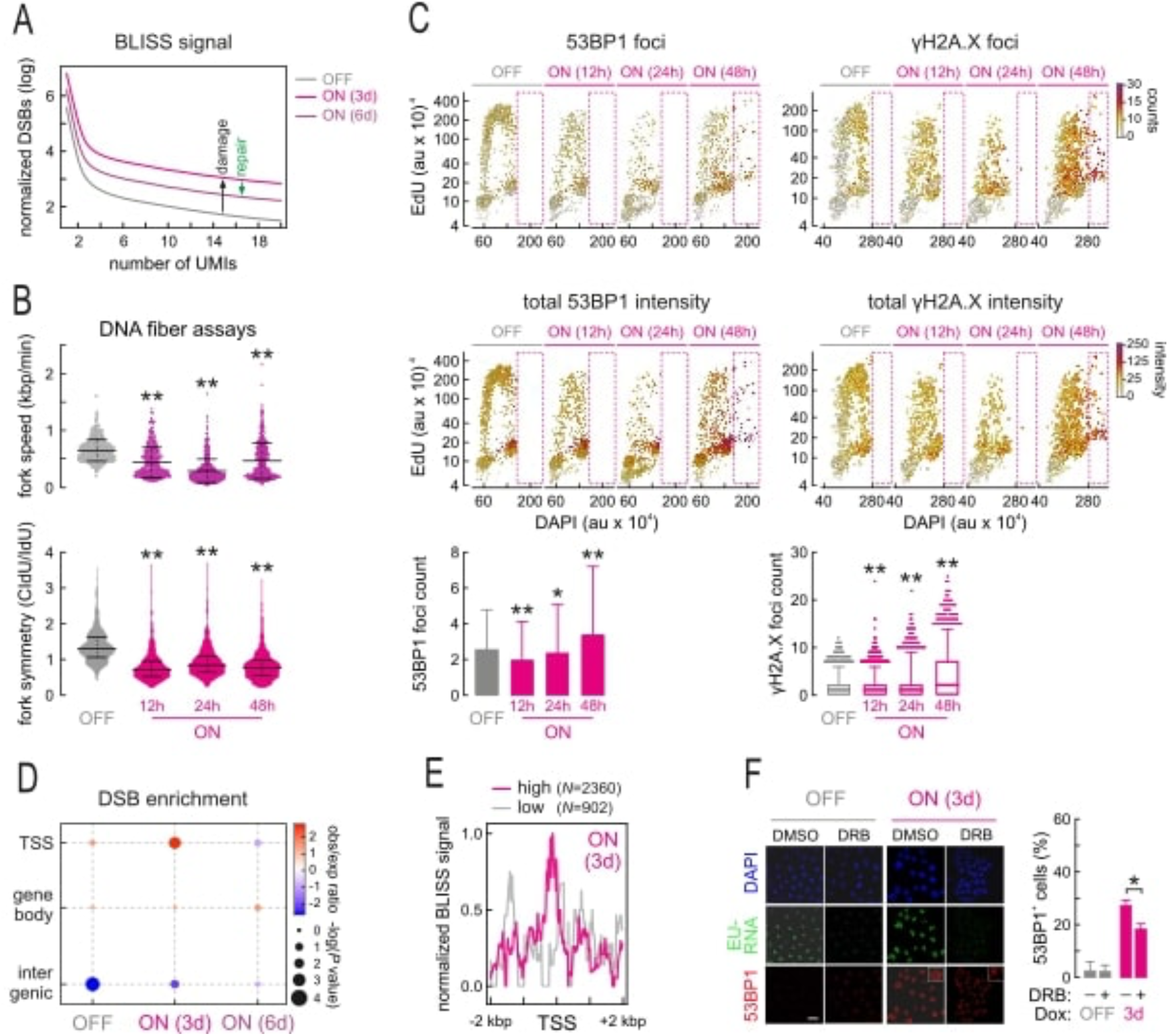
CDC6 induces DNA double strand breaks (DSBs) and alters replication dynamics. (**A**) BLISS data generated at the time points after *CDC6* activation indicated (UMIs: unique molecular identifiers) show strongest DSB accumulation at 3 days followed by 50% reduction at day 6, indicative of DNA repair. (**B**) Violin plots depicting DNA fiber fluorography results show decreased fork rate progression and asymmetry at the time points indicated. **: significantly different to OFF; *P* < 0.01, Student’s t-test (±S.D.; n=3). (**C**) Quantitative image based cytometry (QBIC) of HBECs at the time points indicated shows cell cycle distribution of single cells based on cyclin A and DAPI levels (au: arbitrary units). Foci counts (*top*) and 53BP1 and γH2AX levels (*middle*) are indicated by colour-coding. Bar graphs (*bottom*) show population means (±S.D.) from QBIC measurements. **: significantly different to OFF; *P* < 0.01, Student’s t-test (±S.D.; n=3). (**D**) Dot plot showing increased frequency of DSBs at gene TSSs on the basis of BLISS data. (**E**) Histogram showing BLISS defined DSBs enrichment at gene TSSs upon CDC6 induction. (**F**) Representative immunofluorescence imaging (*left*) of EU-labeled nascent RNA and 53BP1 foci in control HBECs (DMSO) or treated with a transcriptional inhibitor (DRB) at the time points indicated. Bar graphs (*right*) show the percentage (±S.D.; n=3) of cells with 53BP1 foci. *: significantly different to OFF; *P* < 0.05, two-tailed unpaired Student’s t-test.

We suspected that, as a replication factor, deregulated CDC6 would alter replication dynamics and induce “replication stress”, thus explaining DSB formation mechanistically. Indeed, we recorded strong aberrations in the form of reduced fork speed and asymmetry following *CDC6* induction (**Figure 2B**). In addition, the fraction of cells with increased DNA content (>4N) and DNA damage marker expression, indicative of re-replication, progressively increased (**Figure 2C**). Given that DSBs detected by BLISS were particularly enriched at transcription start sites (TSSs) (**Figure 2D**,**E**; in agreement with what was observed by Gothe et al., 2019), we postulated that replication-transcription collisions could occur at these positions. In line with this, global inhibition of transcriptional elongation by RNAPII using DRB significantly reduced the levels of DNA damage response (**Figure 2F**).

Concurrently with DSB emergence, we recorded a prompt (within ∼24 h) and strong increase in RPA foci (**Figure 3A**), a single-strand DNA binding factor and surrogate marker for replication stress [Gorgoulis et al., 2018]. This finding, in combination with our BLISS results, implies that repair predominantly takes place via homologous recombination during S-phase and before the peak of senescence establishment. However, the levels of key components of the main error-free homologous recombination (HR) pathway, that of the synthesis-dependent-strand-annealing (SDSA) repair, like Rad51, BRCA1 and BRCA2, are reduced (**Figure 3B**). In contrast, Rad52 levels and foci are increased upon *CDC6* overexpression (**Figure 3C**,**D**). Thus, in this “BRCAness” environment with low Rad51 levels, DNA repair will predominantly rely on Rad52 activity, which is central to both break-induced-replication (BIR) and single-strand-annealing (SSA) repair routes. Both BIR and SSA are highly error-prone mechanisms contributing to genomic instability and oncogenic transformation [Galanos et al., 2016, 2018; Sotiriou et al., 2016], and we found them significantly activated in ON cells in a Rad52-dependent manner (**Figure 3E**). At the same time, SDSA processivity was strongly reduced, thus satisfying the requirement in our initial hypothesis for a shift from high to low fidelity DSB repair.

**Figure 3.**
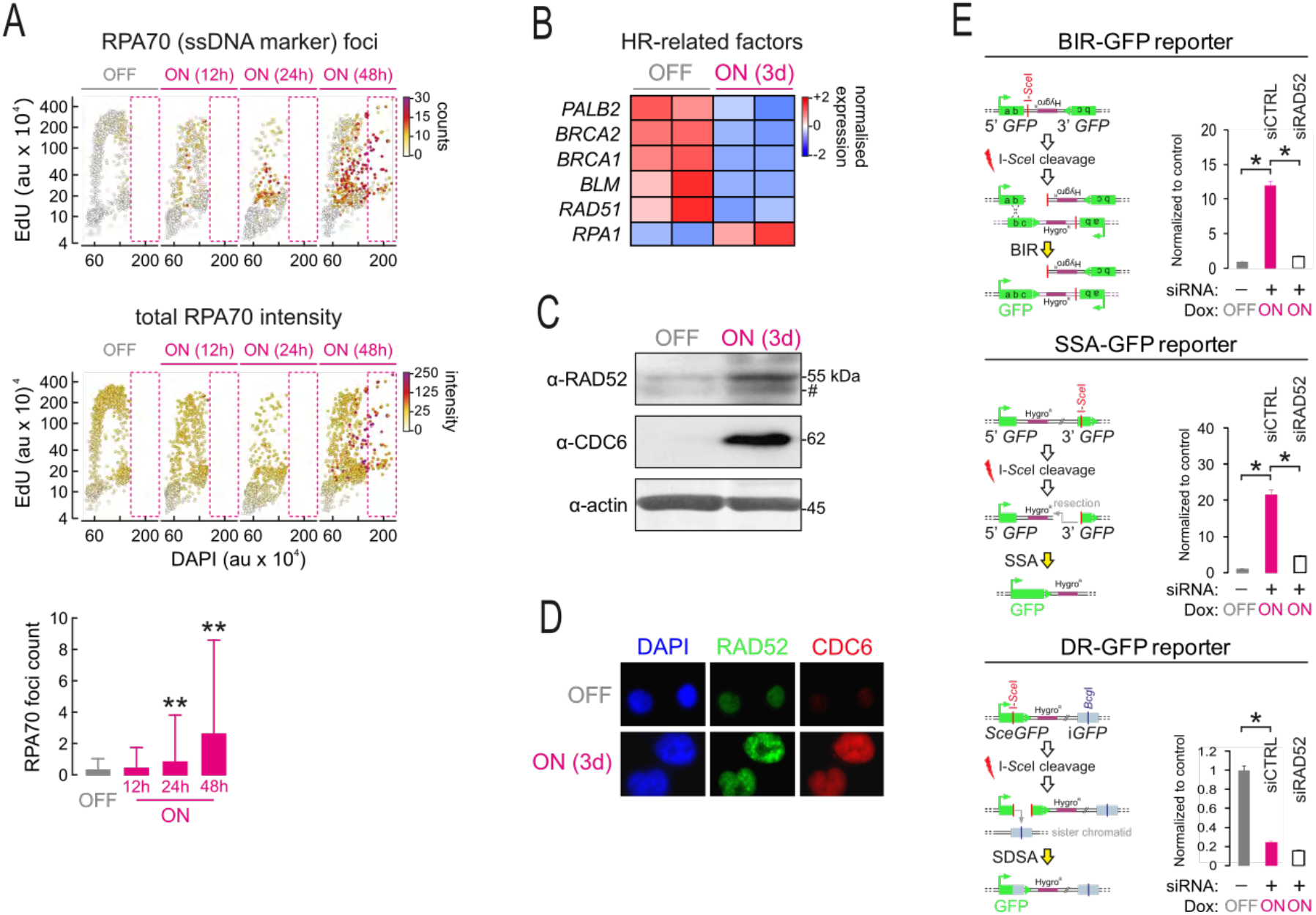
Sustained *CDC6* expression induces replication stress and error-prone DNA repair. (**A**) Quantitative image based cytometry (QBIC) of HBECs at the time points indicated shows cell cycle distribution of single cells based on cyclin A and DAPI levels (au: arbitrary units). Foci counts (*top*) and RPA70 levels (*middle*) are indicated by color-coding. Bar graphs (*bottom*) show population means (±S.D.) from QBIC measurements. **: significantly different to OFF; *P* < 0.01, unpaired two-tailed Student’s t-test (±S.D.; n=3). (**B**) Heatmap showing reduction in the expression levels of the genes involved in error-free homologous recombination (HR) DNA repair upon *CDC6* induction in HBECs (ON). (**C**) Western blots showing RAD52 induction upon *CDC6* overexpression in ON cells. (**D**) As in panel C, but using immunofluorescence imaging of RAD52 and CDC6. (**E**) Reporter assays demonstrating increase (±S.D.; n=3) in RAD52-dependent break-induced replication (BIR; *top*) and in single-strand annealing repair of DSBs (SSA; *middle*). Error-free repair monitored by a synthesis-dependent strand annealing reporter (SDSA; *bottom*) is suppressed. *: *P*<0.05, unpaired two-tailed Student’s t-test.

### “Escape” cells harbor genomic alterations selected early upon senescence entry

Following a senescent period of ∼4 weeks, ESC clones emerged in all three replicates (**Figures S2A-C** and **S3A**). To examine whether traits of DNA damage produced early upon senescence entry are selected and maintained in ESC populations, we employed whole-genome sequencing (WGS). Compared to the uninduced cells, WGS uncovered a broad spectrum of single nucleotide variations (SNVs; sequence alterations of <60 bp) and copy number variations (CNVs; sequence alterations of >60 bp) (**Figures 4A** and **S3B**).

**Figure 4.**
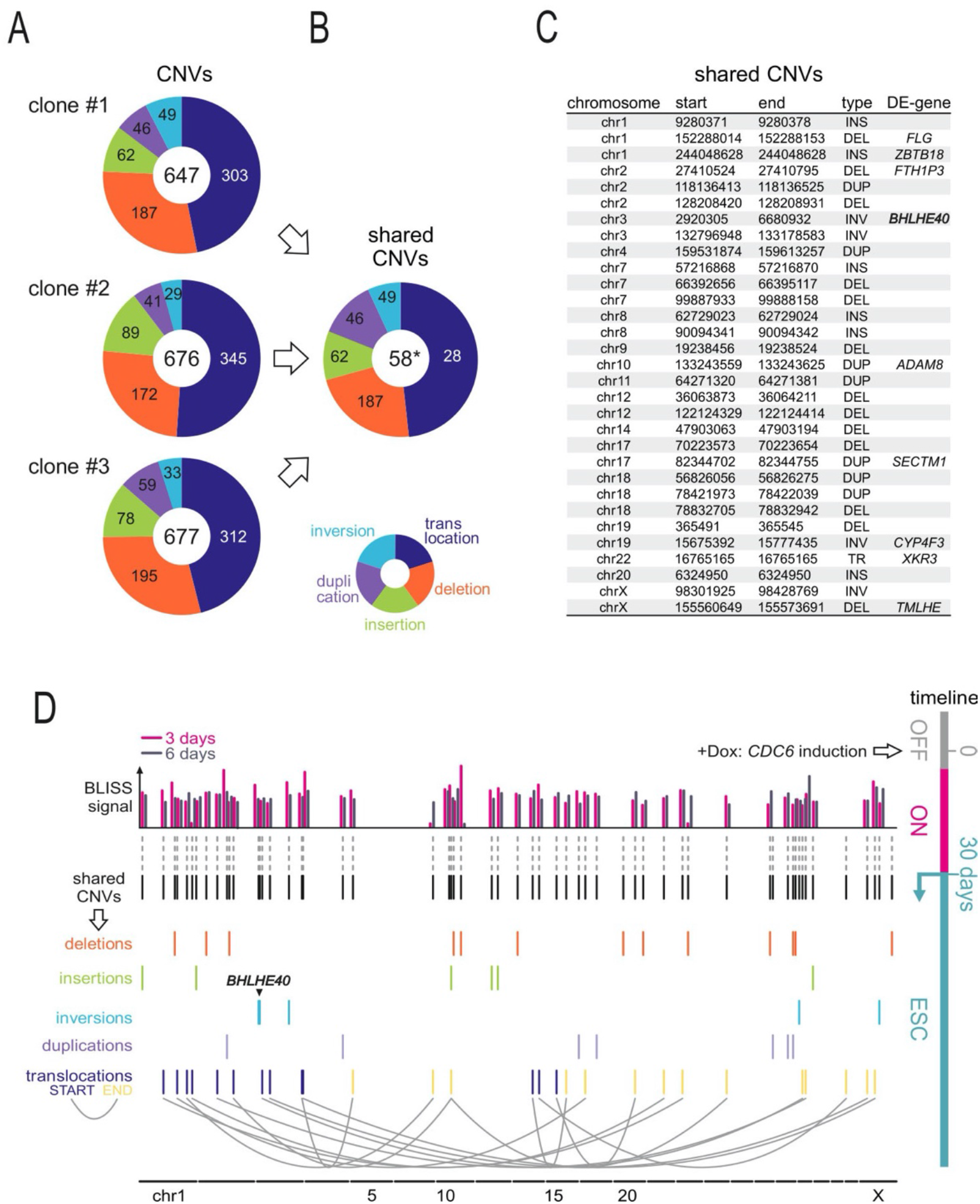
Escape cells harbor recurrent copy number variations (CNVs) aligning to DSBs. (**A**) Pie charts showing the distribution of CNVs identified in each of three independent replicates into five categories. (**B**) Pie charts showing the distribution of the 58 CNVs shared by all the three replicates. *: significantly more than expected by chance; P<0.0001, Super Exact test. (**C**) List of the type and location of all shared CNVs from panel B, alongside any differentially-expressed genes they harbour in ESC cells (*confirmed by RT-qPCR, not in RNA-seq data). (**D**) Superimposing DSB coordinates, as defined by BLISS, with the breakpoints of the shared CNVs from panel B shows overlap in 51 out of the 58 cases.

In more detail, chromosomal distribution of SNVs took a “kategis” form, and we could deduce a mutation signature (**Figure S3C**,**D**) resembling the previously reported “signature 15” associated with mismatch defects seen in stomach and lung cancers [Alexandrov et al., 2013]. Moreover, SNV analysis revealed that our “cancer evolution” model recapitulated two of the most frequently occurring cancer mutations, in *MUC16* and in *NEB* (**Figure S4A-C**), thus validating its relevance. Both mutants associate with poor outcomes in cancer patients [Chugh et al., 2015; Kufe, 2009; Mazzoccoli et al., 2017], with *MUC16* (also known as *CA125*) being an established marker for various cancer types, including lung cancer that is the origin of our cellular system. Although no mutations were found in the *TP53* gene, the most altered gene in cancer (**Figure S4A**), its negative regulator MDM2 increases in “escape” cells leading to its downregulation (**Figure S5A**).

Finally, by interrogating the spectrum of recorded CNVs, we made two observations. First, that — as predicted by our model [Halazonetis et al., 2008; Tsantoulis et al., 2008] — genetic alterations were located within common fragile sites (CFSs; **Table S1**). Second, that 58 out of >650 CNVs per clone were shared by all three replicates (**Figure 4A-C**). Aligning the breakpoints flanking these CNVs to DSB coordinates obtained via BLISS resulted in a striking overlap for 51 out of 58 of them (**Figure 4D**).

In summary, the cancer specific mutational signature (**Figure S3D**), the recapitulation of the *MUC16* and *NEB* mutations seen in patients (**Figure S4**), and the 58 non-random shared CNVs identified in ESC cells (**Figure 4B**,**C**) are all indicative of genomic instability being a decisive determinant in the “escape” from oncogene-induced senescence.

### A large chromosomal inversion uncovers a circadian transcription factor as regulator of “escape”

Given the above data, a fundamental question of our working hypothesis to be addressed is whether genetic alterations obtained early are functionally relevant for the “escape” of the senescence cells that carry them (see **Introduction**). To this end, we noticed a >3.7 Mbp-long heterozygous balanced inversion in the short arm of chr3 in our list of 58 recurring CNVs (**Figures 4B-D, 5A** and **Table S2**). Naturally occurring inversions are generally less susceptible to further recombination, which suggests that genes within such structural variants are selectively “protected” [Wellenreuther and Bernatchez, 2018]. This HBEC-specific inversion encompasses the *BHLHE40* locus (*basic helix-loop-helix family member 40*, also known as *DEC1*) (**Figure 5A**) encoding a transcription factor that belongs to the CLOCK (*circadian locomotor output cycles kaput*) protein family regulating daily circadian rhythm oscillations [Kato et al., 2014; Sato et al, 2016]. Publicly-available ENCODE ChIP-seq data reveal that BHLHE40 exhibits strong and ubiquitous binding across the genome and to regulate >15500 human genes [Rouillard et al., 2016], including many cell cycle regulators (**Figure 5B**).

**Figure 5.**
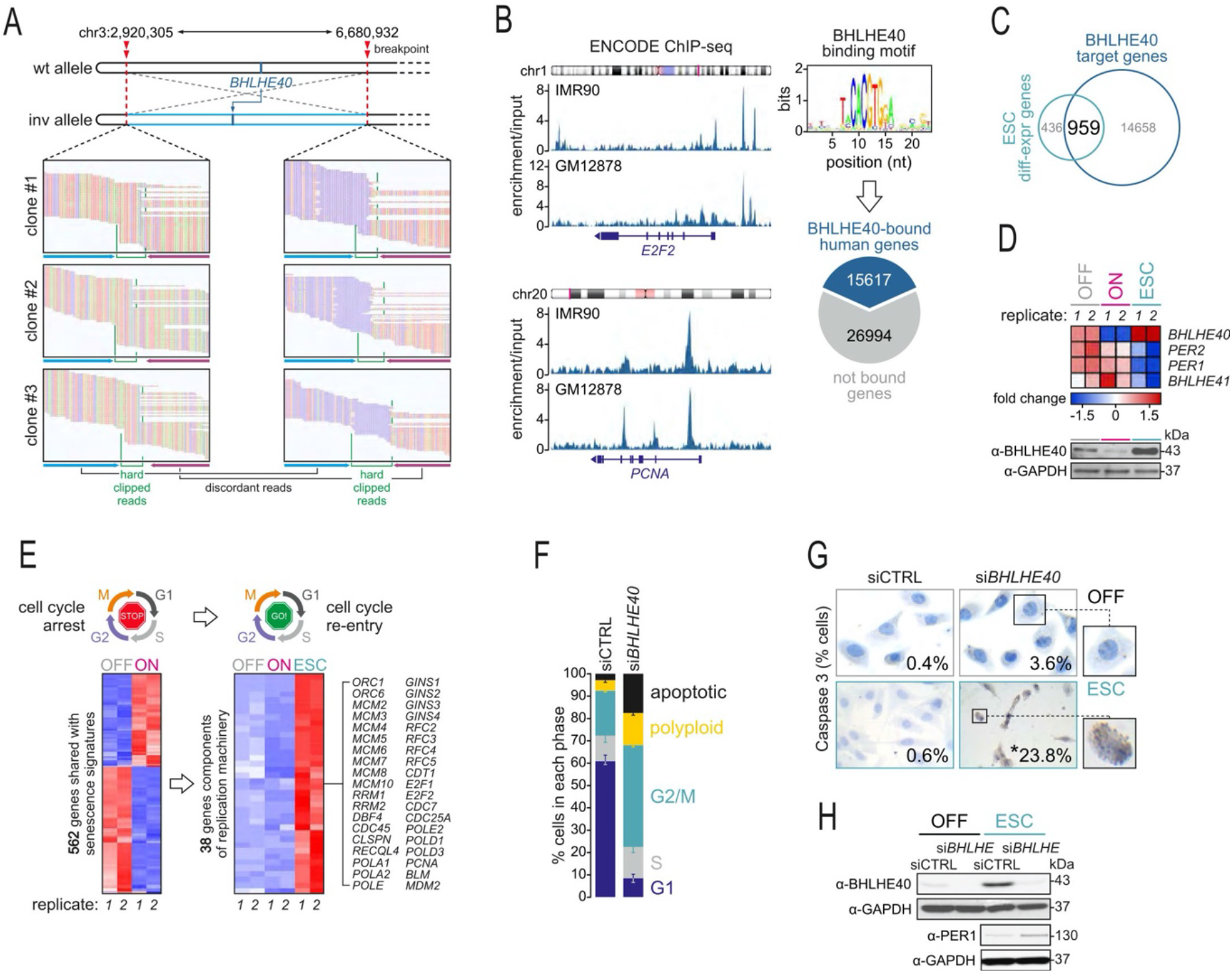
The chr3 inversion harbors *BHLHE40* that is essential for “escape” phenotype maintenance. (**A**) WGS data around the chr3 inversion breakpoints in ESC cells. Hard clipped (*green lines*) and discordantly mapped reads (*blue/purple arrows*) are indicated for all three replicates. (**B**) Representative genome browser views (*left*) of BHLHE40 ENCODE ChIP-seq data from IMR90 and GM12878 cells in the *E2F2* and *PCNA* loci. This data was used to infer the BHLHE40 binding motif logo, and to assign 36.7% of all human genes as its direct targets [Pertea et al., 2018]. (**C**) Venn diagram showing 68.8% of all genes differentially-expressed in ESC cells also being BHLHE40 targets according to ChIP-seq data. (**D**) Heatmap of RNA-seq data shows *BHLHE40*, but not other circadian genes like PER1/2, being selectively upregulated in ESC cells. (**E**) Heatmap (*left*) showing that 25.3% of the 2220 differentially-expressed in ON cells are shared with reported senescence signatures [Hernandez-Segura et al., 2017]. Of these, 38 encode replication machinery components (*right*) and are strongly induced in ESC cells. (**F**) FACS-based cell cycle profiling of control (siCTRL) and *BHLHE40*-knockdown (siBHLHE40) cells showing significantly altered cell cycle progression and increased cell death, denoted with a red arrow (±S.D.; n=3). *significantly more than in control:*P*<0.001, Fisher’s exact test. (**G**) Representative images of control (siCTRL) and *BHLHE40*-knockdown cells (siBHLHE40) immunostained for Caspase-3. Inset numbers indicate the percentage of positive cells (from a minimum of 100 cells counted in each condition). *: *P*<0.01, Fisher’s exact test. (**H**) Western blots showing reciprocal changes in BHLHE40 and PER1 levels upon *BHLHE40*-knockdown in ESC cells, thought to drive apoptosis [Hunt and Sassone-Corsi, 2007].

Notably, >68% of genes differentially-expressed upon “escape” from senescence are reported direct BHLHE40 targets (**Figure 5C** and **Table S3**). Our transcriptome data show that *BHLHE40* is strongly upregulated in ESC cells (also at the protein level; **Figure S5C**), while *PER1/2*, which encode the key circadian factors periodins [Yamada and Miyamoto, 2005; Wood et al., 2009; Kato et al., 2014; Sato et al, 2016], and *BHLHE41* are suppressed (**Figure 5D**). This suggests a direct role for BHLHE40 in the promotion of “escape”. In fact, the circadian circuitry governs, amongst other processes, cell cycle progression and its deregulation affects cell cycle checkpoints and can lead to cancer [Hunt and Sassone-Corsi, 2007; Masri et al., 2013]. Looking into genes encoding replication machinery components, we found 38 key ones that are βoτη strongly reactivated in ESC cells and bound by BHLHE40 (e.g., *BLM, GINS1-4, MCM2-10, PCNA, POLE*; **Figure 5B**,**E**). Among these was also MDM2, the main negative regulator of p53 (**Figures 5E** and **S5A**,**B**).

To test the functional significance of BHLHE40 in this model, we silenced its gene in ESC cells using siRNAs. This led to deregulated cell cycle profiles and increased cell death, as shown via FACS (from 1.89 ± 0.8% cells to 21.25 ± 0.3%; **Figure 5F**) and caspase-3 staining (from 0.5 ± 0.8% cells to 59.9 ± 6.8%; **Figure 5G**). Notably, *BHLHE40* silencing also led to upregulation of PER1 (**Figure 5H**), known to sensitize cells to apoptosis [Gery et al., 2006; Hunt and Sassone-Corsi, 2007]. Together, these results show that *BHLHE40* upregulation is necessary for the maintenance of the “escape” phenotype. The significance of BHLHE40 is also applies to clinical outcomes, as its overexpression is associated with adverse impact on survival in various malignancies, including lung cancer (**Figure S6**).

Apart from the BHLHE40 inversion, which appears to be central in the “escape” phenomenon, a reciprocal translocation involving chromosomes 9 and 22 typically identified in chronic myelogenous leukemia (CML) was also shared by all three ESC populations (**Figure S7**). Finally, all genes laying in the remaining shared CNVs have been associated with the senescence process (for details see **Table S2B**).

### A CRISPR-generated inversion in chr3 suffices for senescence bypass

Subsequently, we tested whether genetic alterations, obtained early upon senescence entry and maintained in ESC cells, are functionally relevant to this transition. In other words, does the inversion in chr3 facilitate “escape” by promoting *BHLHE40* reinduction in response to oncogenic stimuli? To answer this, we first examined BHLHE40 protein levels along a time course from OFF to ESC cells. Baseline levels in OFF cells are reduced upon *CDC6* induction, but markedly increased in the “escape” state (**Figure S5C**). Interestingly, BHLHE40 suppression was partially alleviated by day 6 (**Figure S5C**). This coincides with the window of error-prone DSB repair (**Figure 2A**) and, thus, with the presumed acquisition time of the chr3 inversion.

Next, we used CRISPR-Cas9 editing in HBECs to target sequences within 72 bp (from 2,920,305) and 50 bp (from 6,680,932) of the inversion breakpoints discovered via WGS. We generated two independent clones carrying this 3.7-Mbp heterozygous inversion (**Figures 6A** and **S8A**). “Inverted” cells demonstrated a loss of epithelial features with accentuated spindle morphology, low E-cadherin and emergent Vimentin expression (**Figure 6B**), reminiscent of the metastable state characterizing cells undergoing trans-differentiation [Nieto et al, 2016]. Strikingly, and in accordance to our hypothesis, upon *CDC6* induction the clones carrying this inversion never ceased to proliferate nor did they acquire morphological features of senescence, supporting the notion that they bypass the senescent barrier (**Figures 6C**,**D** and **S8B**,**C**). Notably, at the initial phases of *CDC6* induction, the low S-phase cell percentages observed can be attributed to the particularly energy-demanding state of this metastable phenotype [Nieto et al., 2016] and/or to DDR activation (**Figure S8D**,**E**). This is, nevertheless, not adequate for triggering senescence in this specific “genetic terrain” (**Figure 6B-D**). Soon after this “slow growth” phase (**Figures 6C** and **S8C**), inverted cells do progressively increase their growth rate and invasion capacity (**Figure 6E**,**F**). Critically, both inverted clones overexpressed BHLHE40 upon *CDC6* induction (**Figures 6G**,**H** and **S8F**) and this overexpression drives gene expression changes that favor senescence suppression and cell cycle reentry (**Figure 6I**). In summary, a single inversion in one of the alleles harboring *BHLHE40* suffices for driving constitutive expression of this circadian transcription factor in response to oncogenic overexpression and “escape” from senescence.

**Figure 6.**
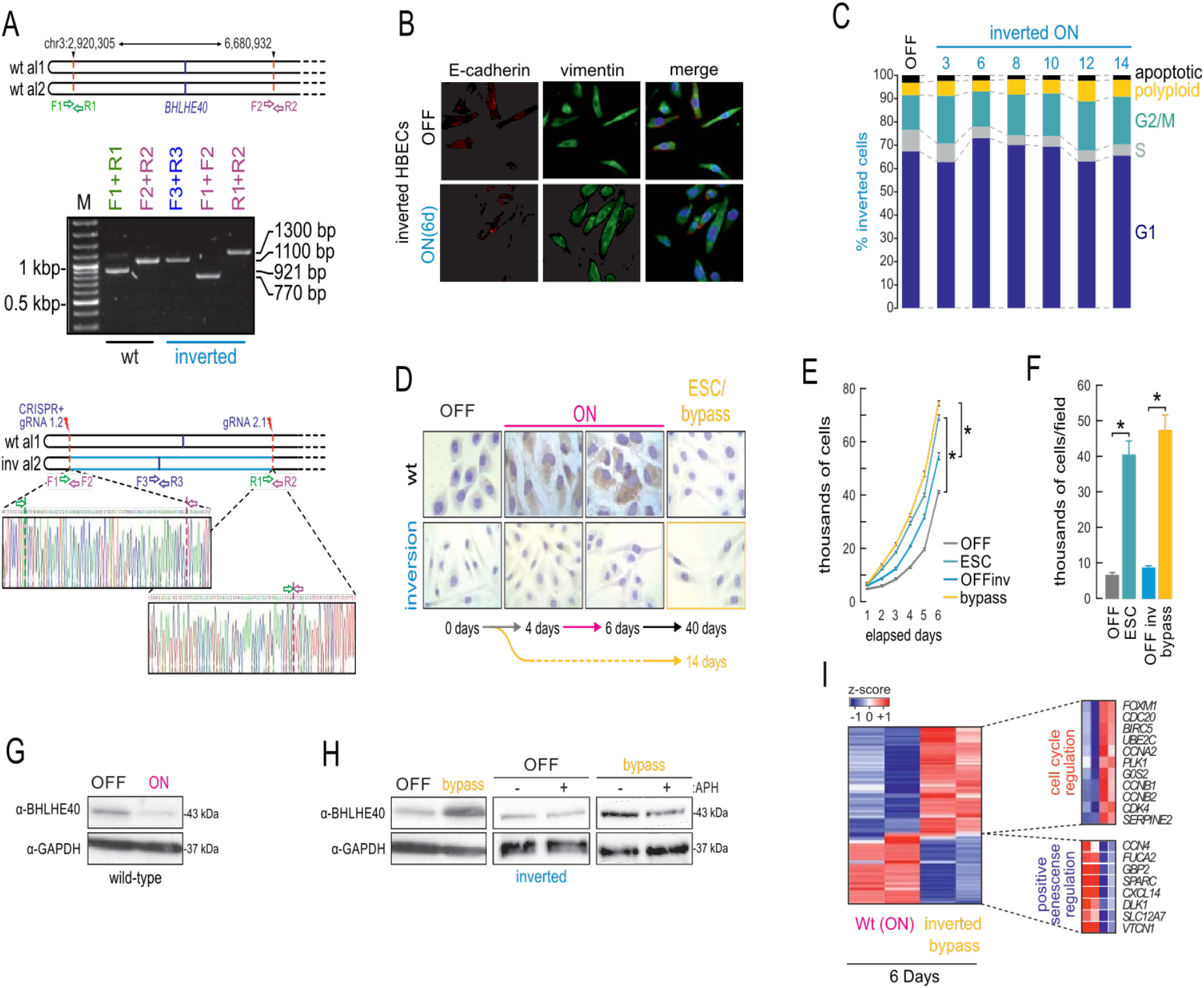
The 3.7-Mbp inversion in chr3 suffices for bypassing CDC6-induced senescence. (**A**) PCR and Sanger sequencing validation of a CRISPR-generated 3.7-Mbp heterozygous inversion in chr3 that closely mimics that discovered in ESC cells using WGS. (**B**) Immunodetection of epithelial (E-cadherin) and mesenchymal markers (vimentin) in “inverted” OFF and 6-day ON cells is reminiscent of cells undergoing transdifferentiation. (**C**) FACS-based cell cycle analysis in “inverted” cells at different times after *CDC6* induction (±S.D.; n=3). (**D**) Representative images (*bottom*) of OFF, ON, and ESC or “bypass” cells stained with SenTraGor to assess senescence-bypass in “inverted” compared to wild-type cells. (**E**) Plots depicting mean proliferation (±S.D.; n=3) in the different states of wild-type and “inverted” cells.*: significantly different to OFF; *P*<0.05, unpaired two-tailed Student’s t-test. (**F**) As in panel E, but quantifying cell invasion capacity (±S.D.; n=3). *: significantly different to OFF; *P*<0.05, unpaired two-tailed Student’s t-test. (**G**) Western blots showing BHLHE40 suppression upon *CDC6*-induction in wild-type cells. GAPDH provides a loading control. (**H**) *Left*: As in panel G, but showing strong BHLHE40 re-expression upon *CDC6*-induction in cells carrying the CRISPR-generated inversion. *Middle/right*: Blots showing aphidicolin (APH) treatment suppresses CDC6-driven BHLHE40 re-expression in “inverted” bypass cells. GAPDH provides a loading control. (**I**) Heatmap of gene expression data depicting inverse patterns for cell cycle and senescence regulators between 6-day *CDC6*-ON wild-type and bypass “inverted” cells.

### Genomic instability-mediated chromatin refolding underlies BHLHE40 induction and “escape” from senescence

It is now understood that changes in three-dimensional (3D) chromosome architecture, like those caused by inversions, may mechanistically explain disease manifestation, including cancer [Ibrahim and Mundlos, 2020]. To test whether this also can also explain *BHLHE40* upregulation, we investigated 3D reorganization in the extended *BHLHE40* locus. We used our “inverted” HBECs to generate high-resolution Hi-C maps from OFF and “senescence-bypass” cells (**Table S4**). Genome-wide comparison of this data revealed that “bypass” cells exhibit an increase in sub-Mbp interactions (**Figure 7A**), accompanied by changes in the identity of compartments. Approximately 10% of A- or B-compartments switch to B or A, respectively, and this switching explains a considerable fraction (almost 50%) of the gene expression changes that underlie senescence bypass (**Figure 7B**). However, only marginal changes to “topologically-associating domain” positions (TADs; Beagan and Philips-Cremins, 2020) were found (**Figure 7C**). These effects are, for the most part, the converse of what was observed for cells transitioning into oncogene-induced or “deep” senescence [Chandra et al., 2015; Criscione et al., 2016].

**Figure 7.**
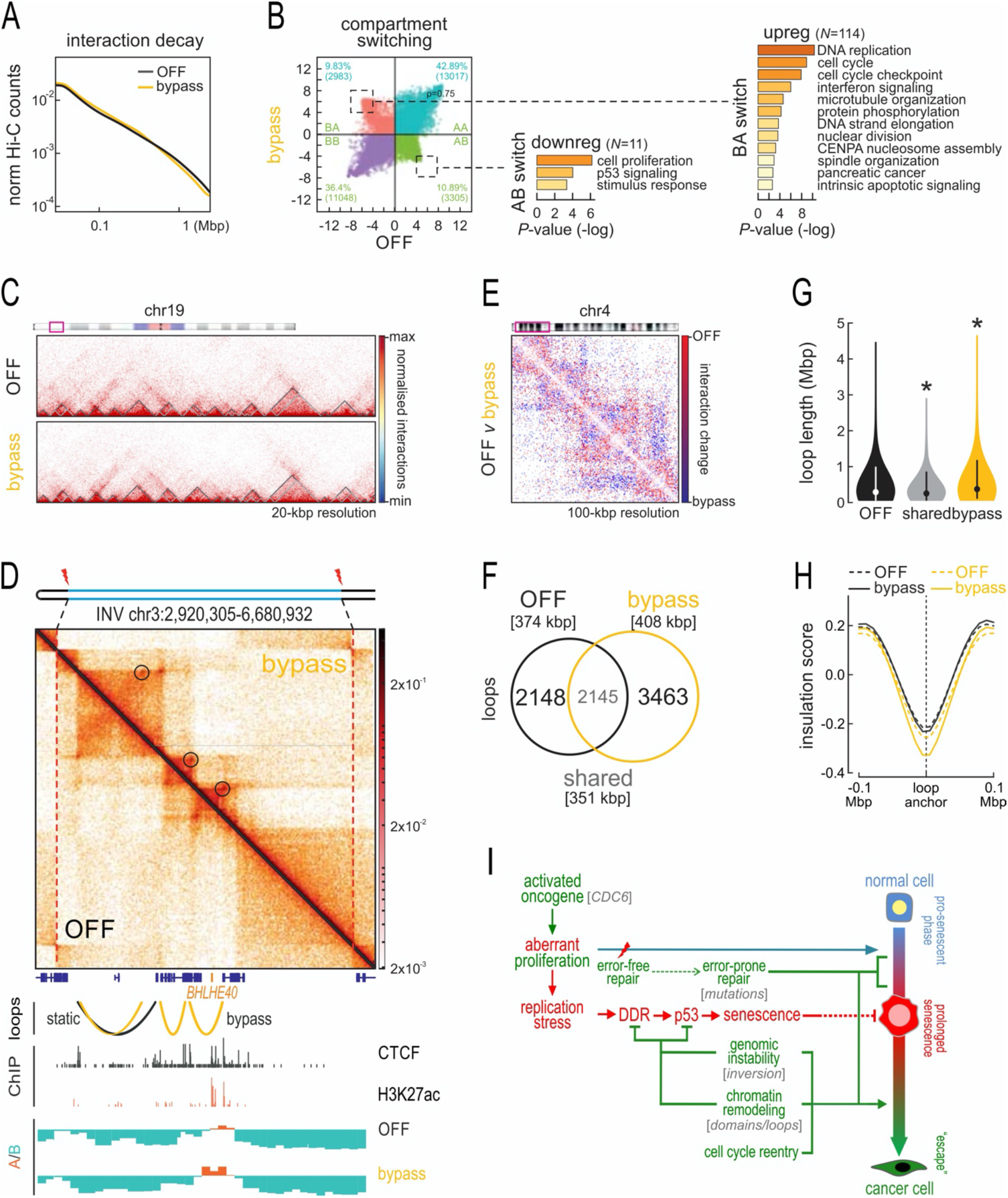
Analysis of spatial chromatin interactions in “inverted” OFF and bypass cells. (**A**) Line plot showing mean interaction strength decay (Hi-C counts) in relation to increasing separation of interacting fragments in OFF (*black*) and bypass “inverted” cells (*yellow*). (**B**) Changes in A/B-compartments in bypass versus OFF Hi-C data. Strong B-to-A and A-to-B switching (*dotted squares*) are indicated, and the GO terms associated with differentially-expressed genes embedded in each switched domain. (**C**) Exemplary Hi-C heatmaps from OFF and bypass cells showing negligible changes in TAD positions for a subregion on chr19. (**D**) Composite Hi-C heatmap depicting interactions from OFF (*bottom*) and bypass “inverted” cells (*top*) in the region harboring *BHLHE40* on chr3. The data is aligned to CTCF and H3K27ac ChIP-seq data from normal OFF HBECs, as well as to A/B-compartment positions from OFF and bypass cells. CTCF-anchored loops emerging upon senescence bypass are denoted on the Hi-C map (*circles*) and aligned below (*yellow arches*). (**E**) Subtracted Hi-C heatmap showing changes in interactions upon transition from OFF to bypass “inverted” cells for a subregion on chr4. (**F**) Venn diagram showing the number of loops unique to OFF and bypass “inverted” cells or shared. Median loop lengths (*square brackets*) are indicated. (**G**) Violin plots showing distribution of lengths for the loops from panel H. *: significantly different to OFF; *P*-value <0.05, Wilcoxon-Mann-Whitney test. (**H**) Line plots showing mean insulation of chromatin interactions in the 200 kbp around loop anchors unique to OFF (*black*) or bypass “inverted” loops (*yellow*) using Hi-C data from OFF (*dotted lines*) and bypass cells (*solid lines*). (**I**) Update on the DNA damage model for cancer development [Halazonetis et al., 2008]. Cells respond to oncogenic stimuli by eliciting senescence as an anti-tumor barrier. The high DNA damage (DSBs) burden amassing during senescence engages error-prone repair mechanisms. Consequently, genetic aberrations accumulate with concurrent chromatin remodeling that provide a “pool” of genomic defects, from which those that facilitate “escape” from senescence, cell cycle re-entry and aggressive features are selected and maintained.

Looking specifically into the 3D organization of chromatin around the inversion region on chr3, we can make three key observations. First, that *BHLHE40* resides in one of the two centrally-located TADs of this extended locus, the long-range contacts of which do not change between OFF and “bypass” cells (**Figure 7D**). Thus, we can rule out the “classical” scenario of *BHLHE40* re-expression due to ectopic contacts with enhancers in adjacent TADs [Ibrahim and Mundlos, 2020]. Second, we recorded the emergence of new loops in this 4-Mbp region, which contribute to the enhanced insulation of the two central TADs from one another (**Figure 7D**, *circles*). Strikingly, a survey of this same 4-Mbp region encompassing *BHLHE40* in publicly-available Hi-C data showed that these two centrally-located TADs appear fused in normal tissue (**Figure S9A**), but well-insulated in cancer cells (**Figure S9B**), thus reflecting our OFF and “bypass” data, respectively. Third, we found that strong loop emergence coincided with the strengthening and broadening of the small A-compartment harboring BHLHE40, which is in line with its more potent activation **Figure 7D**, *bottom*).

Given these effects in the *BHLHE40* domain, we speculated that changes to CTCF loops genome-wide might explain the changes underlying senescence bypass. Indeed, subtracting OFF from “bypass” Hi-C data revealed new long-range contacts emerging (**Figure 7E**). Across all chromosomes ∼3500 new loops arise, while >2000 specific to OFF cells are lost (**Figure 7F**). In line with our subtracted maps, bypass-specific loops are on average larger than OFF-specific ones (**Figure 7G**). Interestingly, and exactly as in the case of the *BHLHE40* domain, these bypass-specific loops arise at positions of existing insulation that become markedly strengthened. At the same time, insulation at the anchors of OFF-specific loops show little fluctuation (**Figure 7H**). Together, this type of changes suggests rewiring of regulatory gene-enhancer interactions. To cite two characteristic examples, we see the emergence of bypass-specific loops in loci suppressed upon senescence bypass. In both cases, these loops trap the two genes, *RRM2* and *NCAPG* (involved in replication and mitosis, respectively), in between consecutive insulated domains to mediate their downregulation (**Figure S10A**,**B; Table S5**). In contrast, *LAP3* finds itself within an emerging bypass-specific loop and is induced (**Figure S10B**).

Furthermore, given that replication origins in mammalians are not defined by specific sequences but rather by structural chromatin context [Antequera, 2004; Cvetic and Walter, 2005] we reasoned that changes in chromatin segment orientation could additionally reorganize the replication process, and in turn affect gene transcription [Lin et al., 2003; Chen et al., 2019; Fisher and Mechali 2003]. The dependence of transcription on replication (S-phase dependence) has been demonstrated in various developmental procedures [Fisher and Mechali 2003]. This, combined with the fact that replication origins can be activated due to replication stress [Courtot et al., 2018], like that induced by *CDC6* overexpression, prompted us to investigate if BHLHE40 up-regulation is linked to replication. Indeed, treating bypass “inverted” cells with aphidicolin markedly reduced the protein levels of BHLHE40, which was not the case for OFF cells (**Figure 6H**). Likewise, wild-type ESC, but not OFF, cells responded the exact same way to aphidicolin by suppressing BHLHE40 levels (**Figure S8G**). Together, such 3D reorganization events can explain gene expression changes leading to senescence bypass.

## DISCUSSION

Entry into senescence is a generalized physiological stress response and as such it is also triggered by oncogene activation to serve as a tumor suppressing mechanism [Gorgoulis et al., 2019]. Still, as with any form of senescence, if the resulting cells are not removed from their niche in a timely manner, an undesirable pro-tumorigenic facet can arise [Rodier and Campisi, 2011; Muñoz-Espín and Serrano, 2014; Gorgoulis et al., 2018; 2019]. This adverse effect has been attributed to the SASP, the secretory cocktail that senescence cells release into their surroundings to trigger chronic inflammation [Gorgoulis et al., 2019]. However, recent reports by us and other proposed that some senescent cells can “escape” this state of oncogene-induced senescence to initiate malignancy [Galanos et al., 2016; 2018; Komseli et al., 2018; Milanovic et al., 2018], but the molecular mechanisms underlying this “escape” still remain obscure.

Here, we present the first mechanistic evidence of how DNA lesions acquired early upon entry into oncogene-induced senescence can drive this phenomenon of “escape”. We exploit normal human bronchial epithelial cell driven to senescence by overexpressing the *CDC6* oncogene. From the populations of these senescent cells, mesenchymal-like, aggressively proliferating cells eventually emerge within ∼40 days. Thus, we essentially mimic “cancer evolution” to find that (i) forced *CDC6* expression induces DSBs genome-wide as early as 3 days post-senescence entry; (ii) these DSBs are predominantly repaired in an error-prone manner; (iii) poorly repaired lesions are actively selected during this “cancer evolution” time course and appear essential for the establishment and/or maintenance of the “escape” clones (**Figure 7I**).

Large genomic cancer studies have shown that the path to malignancy is not uniquely defined, but rather needs to fulfill particular characteristics that allow for the aggressive and unhindered proliferation capacity cancer cells exhibit [Gorgoulis et al., 2018]. We believe that this also applies to the “escape” from senescence. Indeed, our independent ESC clones display recurrent structural and sequence variants that can linked to their phenotype. For example, the precise recapitulation of frequent cancer mutations in *MUC16* and *NEB*, or the resemblance of the ESC SNV signature to that previously discovered in actual patient tumors [Alexandrov et al., 2013]. Another prerequisite for HBEC “escape”, and for most malignant transformation [Aylon and Oren, 2011], is inactivation of the p53 response [Halazonetis et al., 2008]. This also occurs in our model, not via *CDC6*-dependent mutation of the *TP53* locus itself, but rather indirectly via MDM2 upregulation to shut down p53. This course of events is not confined to the bronchial epithelium, but can be recapitulated in human pancreatic duct epithelial cells (HPDECs) that carry an inducible *CDC6* construct and in which p53 function is inactivated via HPV16-E6 transduction [Ouyang et al., 2000]. This is a relevant cell system because *CDC6* overexpression and senescence are frequently detected in precancerous pancreatic lesions [Myrianthopoulos et al., 2019]. As predicted, following *CDC6* induction, HPDECs follow a trajectory that bypasses senescence (**Figure S11**).

Nevertheless, a prominent and recurrent feature in our ESC clones is the 3.7-Mbp heterozygous inversion on chr3. While essentially all types of structural aberrations have been functionally linked to cancer development [Stratton et al., 2009; Danieli and Papantonis, 2020], inversions appear to confer specific properties as regards their selection. Their predominant heterozygous nature allows for their lower recombination potential and, thus, their selective maintenance so that the genes they harbor function in an advantageous “enhanced” mode [Puig et al., 2015; Wellenreuther and Bernatchez, 2018]. Accordingly, the *BHLHE40* gene harbored in our 3.7-Mbp inversion encodes a circadian transcription factor known for controlling a large number of human genes and a variety of processes, including the cell cycle [Hunt and Sassone-Corsi, 2007; Wood et al., 2009; Kato et al., 2014; Sato et al, 2016]. In our system, control of >70% of differentially-regulated genes in “escape” cells can be attributed to BHLHE40. Despite the fact that its expression has been linked to senescence [Collado et al., 2006; Qian et al., 2008], dependence of this “escape” phenomenon on BHLHE40 can be explained by the following sequence of molecular events. Soon after, within <3 days, erroneous repair establishes an inverted locus where this circadian gene is now responsive to *CDC6* overexpression and markedly upregulated. A major player in this appears to be CTCF and its ability to direct loop formation along chromosomes [Rada-Iglesias et al., 2018; Braccioli and de Wit, 2019]. Remodeling of the *BHLHE40* topological domain via the emergence of *de novo* loops coincides with its activation. The resulting abundance of this potent transcription factor is reminiscent of an oncogenic stimulus that can only exert its pro-tumorigenic potential once relieved of the senescence barrier. Such a mode-of-action would then explain contentious reports showing that BHLHE40 triggers senescence or supports cell proliferation, EMT, tumor formation and poor patient survival [Sato et al., 2016; Yamada and Miyamoto, 2005; Qian et al., 2008]. It can also explain those “escape” - relevant gene expression changes that correlate with loop rewiring, in line with the proposed role for BHLHE40 in regulating CTCF binding genome-wide [Hu et al., 2020].

Taken together, our work suggests that it is in the early phase of oncogene-induced senescence that the “genetic seeds” of the forthcoming malignant transformation are planted in chromosomes (**Figure 7K**). Whether “escape” will always be the inevitable destiny of a subset of cells or whether there are cell-autonomous or non-cell-autonomous factors that can dictate this remains to be elucidated. Nonetheless, these findings can have far-reaching clinical implications: the majority of clinically-used chemotherapeutic agents target proliferating cancer cells, while senescent cells represent a dormant compartment tolerant to traditional therapy. Hence, targeting senescent cells can prove of major clinical importance. In light of the expanding field of senotherapeutics [Zhu et al., 2015; Childs et al., 2015; Gorgoulis et al., 2019; Myrianthopoulos et al., 2019], such consideration will be critical in order to prioritize therapeutic choices.

## SUPPLEMENTAL INFORMATION

This manuscript is accompanied by Supplemental Information containing Methods, Figures S1-S11, Tables S1-7, and Video S1.

## DATA AVAILABILITY

All the Hi-C data generated in this study are available via the NCBI Gene Expression Omnibus repository under accession number GSE163371 (reviewer access token: *kfmxuuaxnklzqd*). All the other data are available via the Sequence Read Archive under bioproject PRJNA685322.

## ACKNOWLEDGEMENTS

We would like to thank Dr. Claus Storgaard Sørensen for constructive discussions and Konstantinos Belogiannis and Dr. Athanassios Kotsinas for their input while preparing the manuscript. We also thank Dr. Ioannis S. Pateras, Dr. Teresa Frisan, and Dr. Anastasios Papanastasiou for valuable technical support to this work. V.G.G. is financially supported by: the European Union’s Horizon 2020 research and innovation program under the Marie Sklodowska-Curie grants agreement no. 722729 (SYNTRAIN); the National Public Investment Program of the Ministry of Development and Investment / General Secretariat for Research and Technology, in the framework of the Flagship Initiative to address SARS-CoV-2 (2020SΣ01300001); the Welfare Foundation for Social & Cultural Sciences (KIKPE), Athens, Greece; H. Pappas donation; grants no. 775 and 3782 from the Hellenic Foundation for Research and Innovation (HFRI); and NKUA-SARG grant 70/3/8916. A.P. is supported by the Deutsche Forschungsgemeinschaft via the Clinical Research Unit KFO5002 project grant (CO 1568/2-1) and the Priority Program SPP2202 (grant PA 2456/11-1). Y.Z. is also supported by the IMPRS “Molecular Biology” program, part of the Göttingen Graduate Center for Neurosciences, Biophysics and Molecular Biosciences (GGNB). P.G. is supported by the Danish Cancer Society (R167-A11068) and the Lundbeck Foundation (R322-2019-2577). JB and his team were supported by the Danish Cancer Society (R204-A12617-B153), the Novo Nordisk Foundation (#16854), the Danish National Research Foundation (project CARD, DNRF 125), the Swedish Research Council (VR-MH 2014-46602-117891-30) and the Cancerfonden (#170176).

## AUTHOR CONTRIBUTIONS

C.Z., P.G., A.A., I.M., N.L., K.E., A.M.M.: cell culture and manipulations, immunoblots, FACS, immunofluorescence analysis, immunocytochemistry, SenTraGor staining, combing assays, PCR, invasion assays, QBIC analysis, 3D cell culture; A.A., Y.Z., E.G.G., K.S.: ChIP-seq, Hi-C, CRISPR/Cas9 editing, RNA-seq; D.E.P.: EU assay; R.M., S.G., N.C.: BLISS; T.K., Y.Z., E.G.G., T.K., A.Po., Ai.Po.: bioinformatic analyses; T.K.: survival analyses; P.G., M.D., K.E., J.B., A.P., V.G.G.: data analysis and interpretation, manuscript preparation; J.B., A.P., V.G.G.: experimental design, supervision and project funding, manuscript writing with input for all co-authors.

