## Supplemental Figs S1-S11 and Methods for "Genomic instability is an early event driving chromatin reorganization and escape from oncogene-induced senescence"

### MATERIALS AND METHODS

#### *Cell lines and treatments*

HBEC-CDC6 Tet-ON cells were maintained in Keratinocyte-Serum-Free Medium (#17005-075, Invitrogen) supplemented with 50μg/ml Bovine Pituitary Extract and 5ng/ml hEGF (#17005-075, Invitrogen) at 37°C and 5% $CO_2$ [Komseli et al., 2018]. CDC6 induction was conducted by treatment of the cell culture with 1 μg/ml doxocycline (DOX) (Applichem). Where applied, 5,6-dichloro-1-β-D-ribofuranosylbenzimidazole (DRB, Calbiochem) was used at a final concentration of 100μM and it was added directly in the growth media for the indicated time periods. The cell line used in this study was not found in the database of commonly misidentified cell lines that is maintained by ICLAC and NCBI Biosample. Its identity has been authenticated by STR profiling and is regularly tested for mycoplasma.

#### *siRNA and plasmids transfections*

For BHLHE40 silencing a cocktail of 3 unique 27mer siRNA duplexes - 2 nmol each (OriGene Technologies, Inc, Cat No SR305619) was employed. siRNA gene silencing was performed as previously described, following also the manufacturer's instructions [Galanos et al., 2016].

#### *Protein extraction, cell fractionation and immunoblot analysis*

Protein extraction was performed as described elsewhere [Galanos et al., 2016]. Thirty micrograms of protein from total extracts per sample were adjusted with Laemmli buffer (Sigma) and loaded on acrylamide/bis-acrylamide gels. Gel electrophoresis, transfer to PVDF membrane (Millipore) and signal development with nitro blue tetrazolium/5-bromo-4-chloro-3-indolylphosphate (NBT/BCIP) solution (Molecular Probes) or chemiluminescence have been described before [Galanos et al., 2016]. Alkaline phosphatase-conjugated anti-mouse or anti-rabbit as well as Horse Radish Peroxidase conjugated anti-mouse and anti-rabbit secondary antibodies (1:1000 dilution) (Cell Signaling) were used.

Primary antibodies utilized were: anti-CDC6 (mouse, Santa Cruz, sc9964, 1:500), anti-BHLHE40 (mouse, Santa Cruz, sc101023, 1:200), anti-RAD52 (mouse, Santa Cruz, sc-365341, 1:100), anti-p53 (mouse, Santa Cruz, DO7, 1:500), anti-MDM2 (mouse, Santa Cruz, SMP14, 1:500), anti-PER1 (rabbit, Abcam, ab136451, 1:500), anti-β-actin (rabbit, Cell Signaling, 4967s, 1:1000), anti-GAPDH (rabbit, Cell Signaling, 14c10, 1:2000).

#### *Immunofluorescence analysis*

Indirect immunofluorescence analysis was performed as previously published [Galanos et al., 2016]. Identification of RAD52 foci parameters have been set as indicated before [Galanos et al., 2018]. Secondary antibodies were Alexa Fluor 488 donkey anti-sheep (Abcam, ab150177, 1:500) and Alexa Fluor 568 goat anti-mouse (Invitrogen, no. A11031, 1:500). Image acquisition of multiple random fields was automated on a DM 6000 CFS Upright Microscope (Confocal Leica TCS SP5 II) or a ScanR screening station (Olympus) and analyzed with ScanR (Olympus) software, or a Zeiss Axiolab fluorescence microscope equipped with a Zeiss Axiocam MRm camera and Achroplan objectives, while image acquisition was performed with AxioVision software 4.7.1. Primary antibodies utilized were: anti-CDC6 (mouse, Santa Cruz, sc9964, 1:500), anti-RAD52 (mouse, Santa Cruz, sc-365341, 1:100 for IF), anti-53BP1 (rabbit polyclonal, Abcam #21083, 1:250), anti-CDH1 (E-cadherin) (rabbit monoclonal, Cell Signaling #3195, 1:100), anti-Vimentin (monoclonal, Sigma V6630, 1:100), anti-H3K27ac

(Active Motif #39133. 1:500), anti-H3K27me3 (Active Motif #39155, 1:500). All analyses were performed in triplicate.

***Immunocytochemistry***

For immunocytochemistry analysis cells were grown on coverslips and fixed with 100% ice-cold methanol or 4% formaldehyde (prepared from paraformaldehyde) for 10 min and store at 4°C until staining was performed. Following, cells were permeabilized with 0,3% Triton X-100 in PBS for 5 min at RT. A 10% fetal bovine serum and 3% bovine serum albumin in PBS solution was used as a blocking buffer for 1 h at RT. Primary antibodies were diluted in blocking buffer and incubated overnight at 4°C. Secondary antibodies were: anti-CDC6 (mouse, Santa Cruz, sc9964, 1:500), Ki-67 (mouse, DAKO, MIB-1, 1:500), caspase 3 (rabbit, Cell Signaling, 9662, 1:500). Nuclear signal was evaluated as a positive one. A minimum of 100 cells were counted at high power optical field (x 400).

***Cell growth analysis***

The growth of cells was monitored as previously described [Liontos et al., 2007].

***3D (organotypic/organoid) culture***

First, airway fibroblasts were embedded in type I collagen, allowing contraction of the gel mimicking the underlying submucosa, as previously described [Sato et al., 2006; Ramirez, et al., 2003]. Subsequently, positively selected HBEC-CDC6 Tet-ON cells were seeded on top of the contracted layer and upon attachment of HBECs on the underlying stroma, the organotypic culture was submerged into Keratinocyte-Serum-Free Medium (#17005-075, Invitrogen) supplemented with 50μg/ml Bovine Pituitary Extract and 5ng/ml hEGF (#17005-075, Invitrogen) and then lifted to an air-liquid interface, while cell growth was performed at 37$^o$C with 5% $CO_2$. Following, CDC6 induction was performed as per the 2D culture medium. Finally, matrigels were collected at 6 and 30 days post-induction, formalin fixed and paraffin embedded. Sections were obtained and processed for hematoxylin-eosin and GL13 staining and immunohistochemical analysis as described in previous section.

***Senescence detection with SenTraGor***

SenTraGor$^{TM}$ (available from Lab Supplies Scientific) staining for senescent cells detection was performed as previously described [Evangelou et al., 2017].

***Invasion assay***

Invasion assay was performed as described elsewhere [Sideridou et a.l, 2011; Galanos et al., 2016]. Data from three independent measurements were averaged, and the corresponding SDs were also reported.

***Flow cytometry analysis (FACS)***

Cell cycle analysis was determined using a FACS Calibur (Becton-Dickinson) as previously before [Galanos et al., 2016].

***5'-EU incorporation based nascent RNA assay***

*In situ* detection of nascent RNA was performed with the Click-iT Alexa Fluor 488 Imaging Kit (Invitrogen, Molecular Probes) as described elsewhere [Komseli et al., 2018].

***QIBC analysis***

Quantitative image-based cytometry (QIBC) analysis was performed essentially as previously described [Ochts et al., 2016]. In brief, images were taken with a ScanR inverted microscope High-content Screening Station (Olympus) that was equipped with wide-field optics, a 20×, 0.75-NA (UPLSAPO 20×) dry objective, fast excitation and emission filter-wheel devices for DAPI, FITC, Cy3, and Cy5 wavelengths, an MT20 illumination system, and a digital monochrome Hamamatsu ORCA-R2 CCD camera. Images were obtained in an automated fashion with the ScanR acquisition software (Olympus, 2.6.1). Depending on cell confluency, 25 to 49 images were acquired containing at least 1,000 cells per condition. Acquisition times for the different channels were adjusted for non-saturated conditions in 12-bit dynamic range, and identical settings were applied to all the samples within one experiment. Images were processed and analyzed with ScanR analysis software. First, a dynamic background correction was applied to all images. The DAPI signal was then used for the generation of an intensity-threshold-based mask to identify individual nuclei as main objects. This mask was then applied to analyze pixel intensities in different channels for each individual nucleus. For analysis of IR-induced foci, additional masks were generated by segmentation of the respective images into individual spots with intensity-based or spot-detector modules included in the software. Each focus was thereby defined as a sub-object, and this mask was used for quantification of pixel intensities in foci. After this segmentation of objects and sub-objects, the desired parameters for the different nuclei or foci were quantified, with single parameters (mean and total intensities, area, foci count, and foci intensities) as well as calculated parameters (sum of foci intensity per nucleus). These values were then exported and analyzed with TIBCO Software, version 5.0.0. This software was used to quantify absolute, median, and average values in cell populations and to generate all color-coded scatter plots. Within one experiment, similar cell numbers were compared for the different conditions (at least 1,000 cells), and for visualization low x-axis jittering was applied (random displacement along the x axis) to make overlapping markers visible.

***DR-GFP, SA-GFP and BIR-GFP reporter assays***

HBEC-CDC6 Tet-ON cells were transiently transfected with the GFP based reporter constructs for synthesis-dependent strand annealing (DR-GFP), single strand annealing (SA-GFP) and break induced replication (BIR-GFP), as previously described [Galanos et al., 2018]. To monitor repair of I-*Sce*I- generated DSBs, cells were transiently cotransfected with 1 μg of the I-*Sce*I expression vector HA-ISceID44A (Addgene #59424) using the Effectene reagent (Qiagen). DSB repair efficiency upon CDC6induction was determined by quantifying GFP-positive cells via flow cytometry FACS Calibur (Becton Dickinson) 48h after transfection.

#### *DNA fiber fluorography (combing assay)*

The assay was conducted as previously described [Galanos et al., 2016]. Briefly, HBEC-CDC6 Tet-ON cells were grown in the presence or absence of doxycyclin for the indicated time points (see **Figure 2**) and then pulsed-labeled with 25μM CldU for 20min, and then labelled with 250μM IdU for 20min. Cells were then harvested and lysed on glass slides in spreading buffer, DNA was denatured and stained using rat anti-BrdU/CldU (1:1000, OBT0030F, Immunologicals Direct) and mouse anti-IdU/BrdU (1:500, clone B44, Becton Dickinson) antibodies.

#### *Breaks Labeling In Situ and Sequencing (BLISS)*

"Breaks Labeling In Situ and Sequencing" (BLISS) analysis was performed as previously described in [Yan et al., 2017; Bouwman et al., 2020]. Briefly, the method consists of following main steps: i) upon harvesting of cells from multi-well plates, approx. 2 million cells were fixed in suspension with 4% formaldehyde for 10 min at room temperature, ii) DSBs ends were *in situ* blunted, iii) next they were tagged with dsDNA adapters containing sample barcodes, UMIS (unique molecular identifiers), RA5 adapter and T7 promoter, iv) tagged DSB ends were linearly amplified using *in vitro* transcription and v) the resulting RNA was used for library preparation and sequencing. BLISS data were analyzed as described below.

#### *Next Generation Sequencing and Bioinformatics analysis*

For whole-genome sequencing (WGS), library preparations were as described previously [Galanos et al., 2018]. SAMtools *mpileup* and *bcftools* [Li et al 2009]*, GATK tools*, the GATK source bundle and the GATK best practices guide [Van der Auwera et al, 2013], were used for identification and filtering of the SNPs and INDELs. Variations that were unique in the "escaped" cells were normalized based on the sequencing depth of each experiment. Copy number and structural variants were determined using MANTA [Chen et al., 2016] and annotated on the Human reference genome using ANNOVAR [Wang et al., 2010]. For BLISS data, DNA Double Stranded Breaks (DSBs) were normalized for total mapped reads and for the total number of used cells for each replicate. The aggregation of Unique Molecule Identifiers (UMIs) and the frequency of DSBs in various genomic regions were calculated using in-house R scripts (available on request).

#### *RNA isolation, sequencing, and data analysis*

6-day ON and senescence-bypass "inverted" HBECs were harvested in Trizol (Invitrogen) and total RNA was isolated and DNase-treated using the Direct-zol RNA miniprep kit (Zymo Research) as per manufacturer's instructions. cDNA libraries were next generated using the TruSeq RNA library kit (Illumina) via selection on poly(dT) beads. The resulting libraries were single-end sequenced to >50 million reads on a HiSeq4000 platform (Illumina). Raw reads were mapped to the human genome (hg19) using STAR aligner (version 2.5.3a) [Dobin et al, 2013]. Samtools (version 0.1.19) [Li et al, 2009] were used for data filtering and file format conversion, while HTseq count (version 0.5.4p3.) algorithm [Anders et al, 2015] was used to assign aligned reads to exons using the following command line «htseq-count –s no –m intersection -nonempty». Normalization of reads and removal of unwanted variation was performed with RUVseq [Risso et al., 2014]. Differential gene expression was computed using DESeq [Anders, 2010], and significantly deregulated genes (fold change cut-off 1.5 and P-value≤0.05) are listed in **Table S6**.

#### *Chromatin immunoprecipitation (ChIP), sequencing, and data analysis*

ChIP was performed on 10-15 million cells crosslinked in 1% PFA/PBS at RT for 10 min, and quenched in 0.125M ice-cold glycine. ChIP material was prepared as previously described [Ford et al., 2014], and sonication was performed using a Bioruptor sonicator and adjusting fragment size to 200-500 bp. For the IP the following polyclonal antisera were used: anti-CTCF (61311, Active Motif) and anti-H3K27ac (39133, Active Motif). ChIP-seq libraries were sequenced on a HiSeq4000 platform (Illumina) to at least 25 million reads per sample, and data was analyzed according to ENCODE guidelines. In brief, reads were aligned to the reference human genome (GRCh37/hg19) using BWA-MEM [Li and Durbin, 2010], and *SAMtools* [Li et al., 2009] was used to convert .SAM into .BAM files. Peaks were defined using MACS2 (ver. 2.1.2) [Zhang et al., 2008] and a self-defined model further refined by first shifting each paired mapped read by 100 bp towards its 3' end and then extending each tag position toward its summit, before the default deduplication option of MACS2 was applied to remove PCR artifacts. Finally, maximum tags at each location were calculated on the basis of a binomial distribution with a *P*-value cutoff of $10^{-6}$; in the case of two or more peaks overlapping, the one with the lowest *P*-value was kept while the others were discarded.

#### *Genome-wide chromosome conformation capture (Hi-C) and data analysis*

*In situ* Hi-C on HBECs of different states and genotypes was performed and controlled for quality using the Arima Hi-C kit as per manufacturer's instructions. All resulting libraries that met the QC criteria set by the manufacturer were paired-end sequenced on a NovoSeq6000 platform (Illumina) to at least 0.5 billion reads. For data analysis, reads were mapped to the reference human genome (GRCh37/hg19) using Bowtie (ver. 23.4.1) [Langmead and Salzberg, 2012] with the "--reorder" flag. Local mapping was used to increase mapping rates due to the inherent presence of chimeric reads. All preprocessing and downstream analysis was performed using HiCExplorer (ver. 3.2) [Ramirez et al., 2018] to remove unmappable reads, non-uniquely mapped reads and low-mapping-quality reads, as well as duplicated pairs (i.e., starting and ending with exactly the same location), dangling-ends (i.e., digested but not ligated), self-circularised (i.e., reads pairing within <25 Kbp and facing outwards), same-fragment (i.e., read pair locating in the same restriction enzyme fragment) or self-ligated reads (i.e., having a restriction site in between the read pair within <800 bp). Next, genome-wide contact matrices were generated in the form of *.cool* files, in which the genome was binned into different sizes (resolution) — 10 kb, 20 kb, 50 kb and 100 kb — for different downstream usage. To facilitate comparison between different samples, all Hi-C interaction counts were normalized and then balanced using the Knight-Ruiz (KR) matrix balancing algorithm [Knight and Ruiz 2012]. Hi-C matrices stored in *.cool* files were visualized using HiGlass [Kerpedjiev et al., 2018] as interactive heatmaps. To make zooming-in and -out possible, normalized and balanced *.cool* files at 10 Kbp resolution were converted to multi-resolution cooler files called *.mcool* files using Cooler [Abdennur and Mirny 2019]. For calling A/B compartments, 100 kbp-resolution and Pearson-transformed matrices were used to calculate the first eigenvector, which was then integrated with own H3K27ac ChIP-seq data to mark A-compartments. TADs were assigned using 20 kbp-resolution matrices using the function embedded in HiCExplorer based on deduced *z*-scores and with a *P*-value cutoff of 0.01. Finally, loops we detected as previously described [Rao et al., 2014] by computing a negative binomial distribution of 10 kbp-resolution Hi-C data and using Anderson-Darling/Wilcoxon rank-sum tests and a *P*-value cutoff of 0.05; loop lengths were restricted to 0.1-2 Mbp (to avoid signal contamination from the diagonal of Hi-C matrices), and compared to CTCF ChIP-seq data to identify loops with CTCF-bound anchors.

***CRISPR/Cas9 inversion generation***

*Design of gRNAs*. Based on the WGS data (see corresponding section), 20-nt sgRNAs were designed around each breakpoint. Two complementary DNA oligos for each sgRNA were annealed generating 5'overhangs consisting of CACC(G) and AAAC. gRNA1 and gRNA2 were chosen due to high specificity and small distance from the exact breakpoints (listed in supplemental material). They were cloned into – Cas9 expression plasmids - pSpCas9(BB)-2A-GFP (PX458) and pU6-(BbsI)_CBh-Cas9-T2A-mCherry, respectively, which had been already digested with BbsI. In this way, sgRNAs were integrated next to the gRNA scaffold of the particular vector (**Table S7**).

*Transfection and FACS sorting*. HBECs were cultured in Keratinocyte (serum free medium) (#17005042) without antibiotics supplemented with 25 mg Bovine Pituitary Extract and 2.5 µg EGF, Human Recombinant. Delivery of 2.5µg from each plasmid, coding for one sgRNA and Cas9, was performed via double transfection of HBECs two days after plating 8x$10^4$ cells per well in a 6-well plate (reaching 80% confluency) with FuGENE ® HD Transfection Reagent (Promega #E2311) (4:1 FuGENE® HD Transfection Reagent: DNA Ratio). FACS sorting of double positive (GFP and mCherry) cells gave rise to a large number of clones, subsequently cultured in 96-well plates.

*DNA extraction and PCR screening.* After harvesting cells from 96-well plates in 30 µl Trypsin/EDTA 1x (#15400054), followed by a neutralization step with an equal volume of Trypsin Neutralizer Solution (#R002100), half of the cells were lysed by adding 30 µl of Lysis Buffer (50 mM KCl, 1 0mM TRIS pH: 8.3, 2.5 mM $MgCl_2$, 0.45% NP40 and 0.45% Tween20) containing Proteinase K (1 µl of 20 µg/µl Proteinase K for every 50 µl of Lysis Buffer), and heating for 45 min at 60°C followed by 10 min at 80°C to inactivate Proteinase K. The other half of the cells were kept in culture. 4µl of the lysate were used as genomic DNA for PCR. Two pairs of forward and reverse primer were designed around each breakpoint (**Table S7**). PCR product of F1/R1 and F2/R2 manifest the wild type genomic DNA, while F1/F2 and R1/R2 give product in case that the area has been inverted. PCR products were submitted for Sanger sequencing verification (see below).

***Sanger sequencing***

PCR products were purified and submitted for Sanger sequencing as previously described [Karakaidos et al., 2004].

***Survival analysis***

Data on survival analysis was obtained from a public database Kaplan-Meier plotter (http://www.kmplot.com) [Nagy et al., 2018], except for breast and prostate cancer data for which a separate Log-rank (Mantel-Cox) survival analysis, with Bonferroni correction, was performed on data retrieved from Metabric and TCGA, respectively.

### SUPPLEMENTAL FIGURES

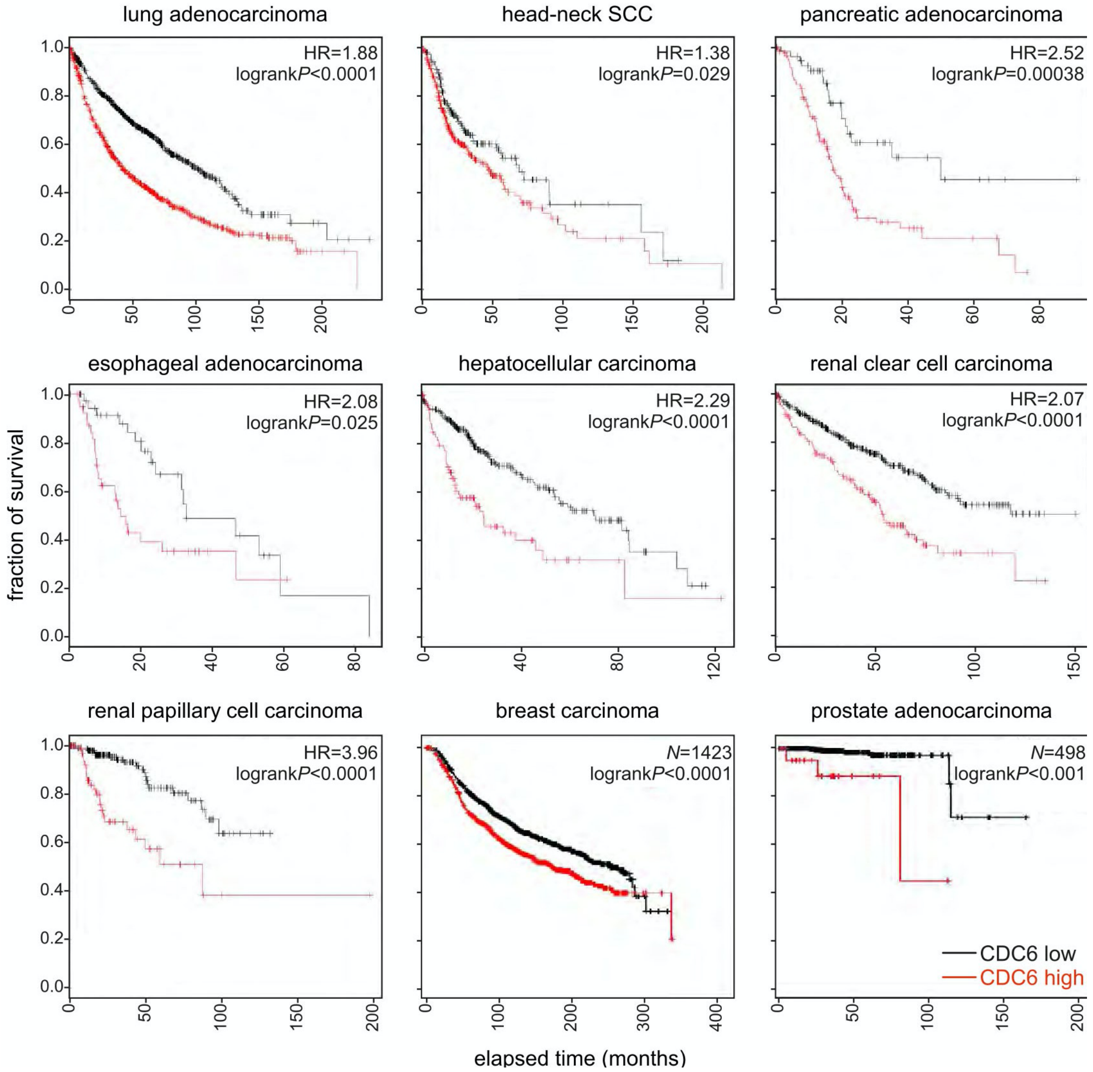


**Figure S1.** ***CDC6*** **overexpression is associated with poor survival of cancer patients.** Kaplan-Meier survival plots generated using public data from tumors stratified as "high" (*red line*) or "low" CDC6-expressing (*black line*; http://www.kmplot.com). Plots for breast and prostate tumors were generated using data from Metabric and TCGA, respectively.

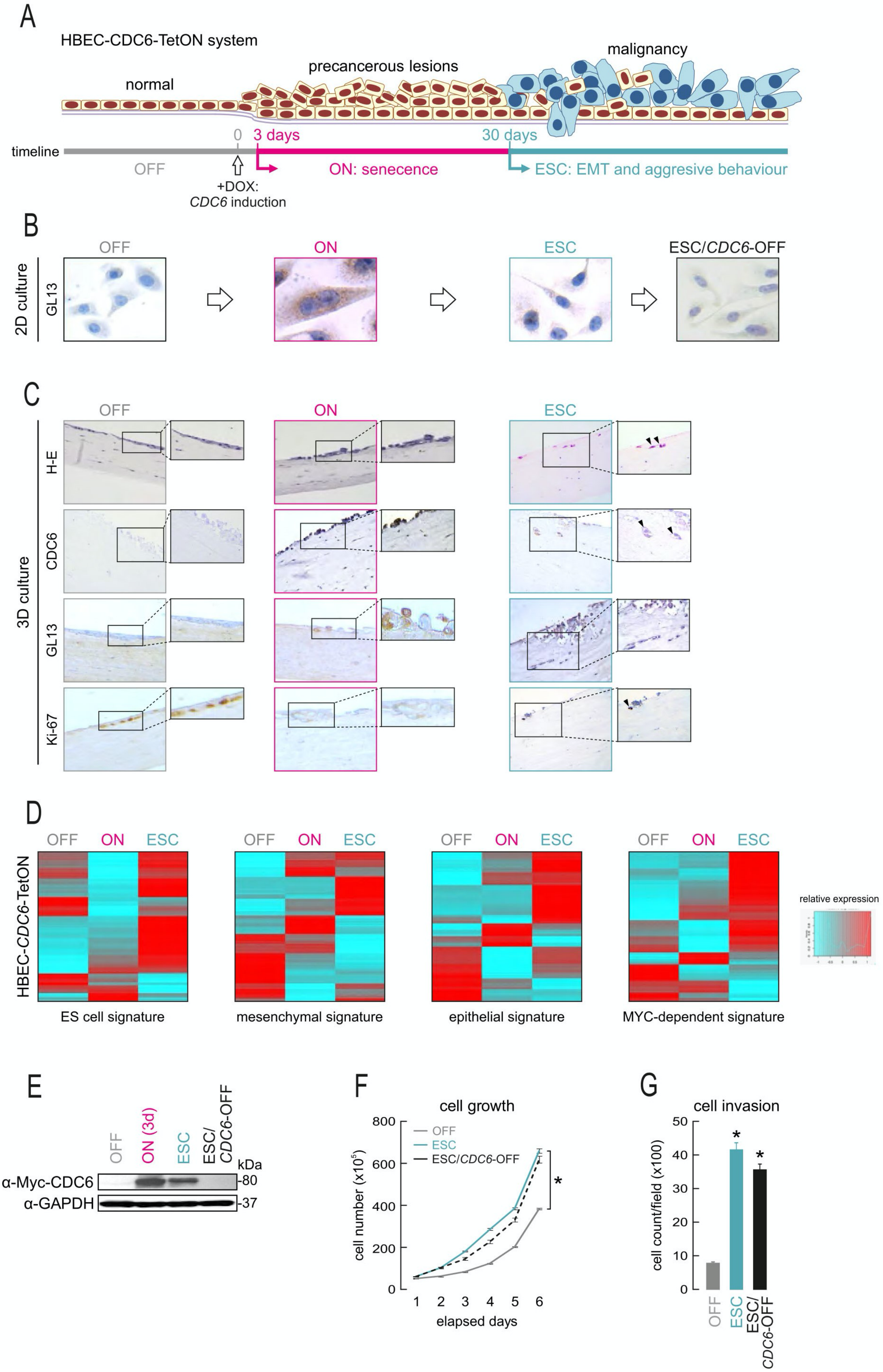
A
HBEC-CDC6-TetON system
normal
precancerous lesions
malignancy
timeline
0
3 days
30 days
OFF
+DOX:
CDC6 induction
ON: senecence
ESC: EMT and aggresive behaviour
B
2D culture
GL13
OFF
ON
ESC
ESC/CDC6-OFF
C
3D culture
H-E
CDC6
GL13
Ki-67
D
HBEC-CDC6-TetON
ES cell signature
mesenchymal signature
epithelial signature
MYC-dependent signature
relative expression
E
ON (3d)
ESC/CDC6-OFF
kDa
α-Myc-CDC6
α-GAPDH
-80
-37
F
cell growth
cell number (x10^5)
elapsed days
G
cell invasion
cell count/field (x100)

311

**Figure S2. *CDC6* induction in HBECs triggers a process recapitulating all stages of malignant transformation.**

(**A**) Overview of our model of cancer evolution based on inducible *CDC6* expression in HBECs.

(**B**) Representative images of HBECs grown in 2D culture and immunostained for GL13 (Sentragor). *CDC6* induction forces cells into senescence (ON). After ~30 days, a small subset of cells “escape” senescence (ESC) to re-enter the cell cycle and adopt an EMT phenotype. Shutting-off *CDC6* in ESC cells (ESC/*CDC6*-OFF) does not reverse this phenotype.

(**C**) Representative images of HBECs grown in 3D organotypic conditions and immunostained for H-E (hematoxylin-eosin), CDC6, GL13 (Sentragor), and Ki-67 following the same timeline as in panel B. Non-induced cells (OFF) recapitulate the upper respiratory epithelium. Upon *CDC6* induction, cells enter senescence and form spheroids. Prolonged *CDC6* induction gives rise to ESC cells with an EMT phenotype (*arrowheads* in H-E-stained ESC cells) and renewed proliferative capacity (*arrowhead* in Ki-67-stained ESC cells) that invade the supporting collagen matrix (*arrowheads* in CDC6-stained ESC cells).

(**D**) Heatmaps showing that ESC cells display a mixed gene expression signature, consisting of embryonic, mesenchymal, epithelial and Myc-dependent markers.

(**E**) Western blots showing changing levels of Dox-induced CDC6 in HBECs.

(**F**) Line plots quantifying sustained proliferation (mean ±S.D.; n=3) of ESC/*CDC6*-OFF cells. *: significantly different to OFF: $P<0.05$, unpaired two-tailed Student’s t-test.

(**G**) As in panel F, but with bar plots quantifying cell invasion capacity.

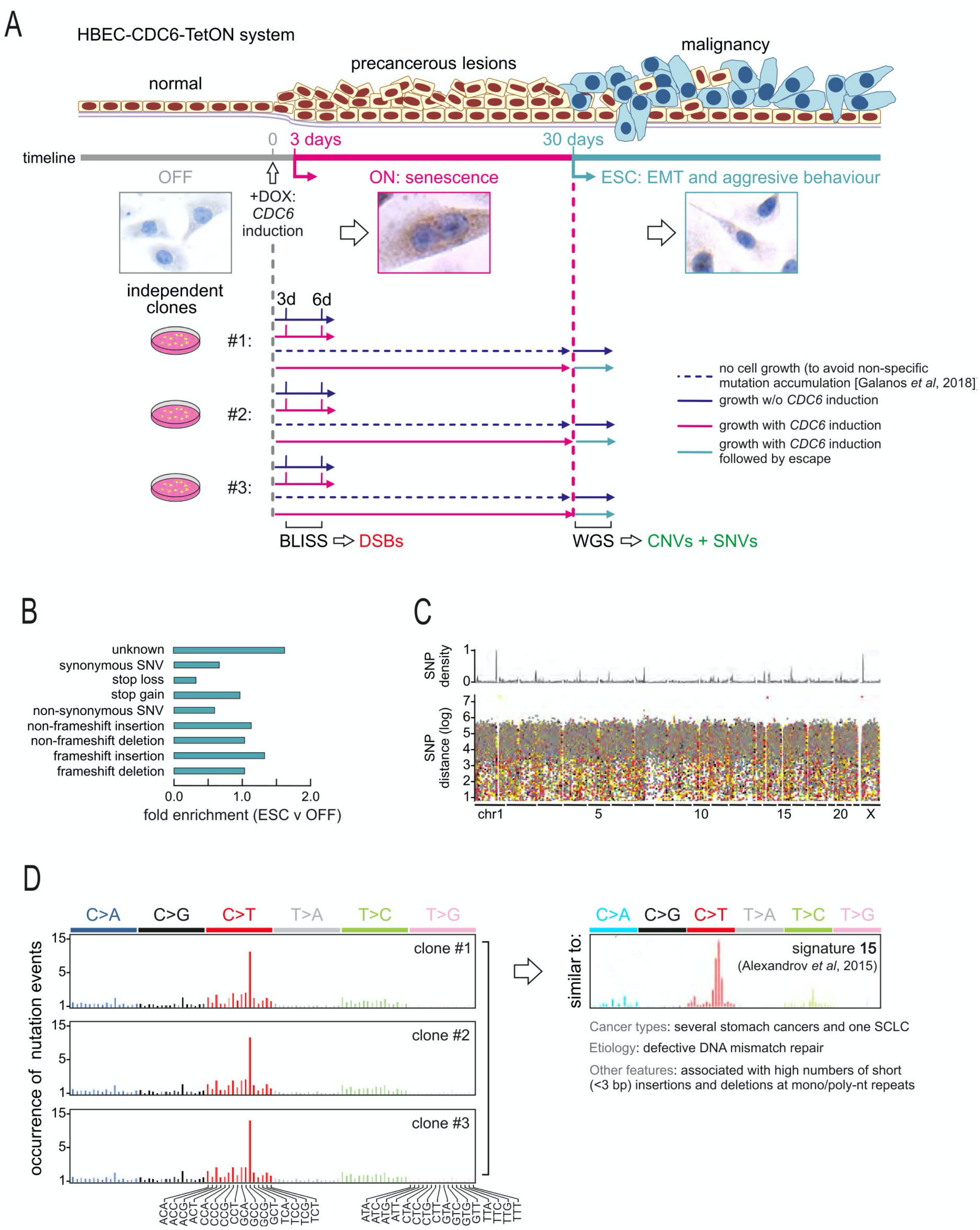


**Figure S3. CDC6-driven mutational landscapes in ESC cells.**
(**A**) Overview of our three independent “escape” experiments. BLISS was applied to identify DSBs occurring after 3 or 6 days of *CDC6* induction. Then, whole-genome Sequencing (WGS) was performed on ESC cells to map genetic alterations in respect to damage that occurred at early time points. OFF cells that served as controls for WGS analysis were only initiated for culture at the time when ESC cells emerged to avoid non-specific accumulation of genetic alterations in the prolonged stationary period of senescent ON cells.
(**B**) Bar plots showing the type and relative enrichment of SNVs in ESC cells using WGS data.

(**C**) WGS-derived SNP density plots aligned to a “kategis” SNV distribution in ESC genomes.

(**D**) Bar plots showing the occurrence of specific SNVs in each of the three independent replicates that represent a *CDC6*-specific mutational signature similar to that previously reported for stomach cancer and one small lung cell carcinoma [Alexandrov et al., 2013].

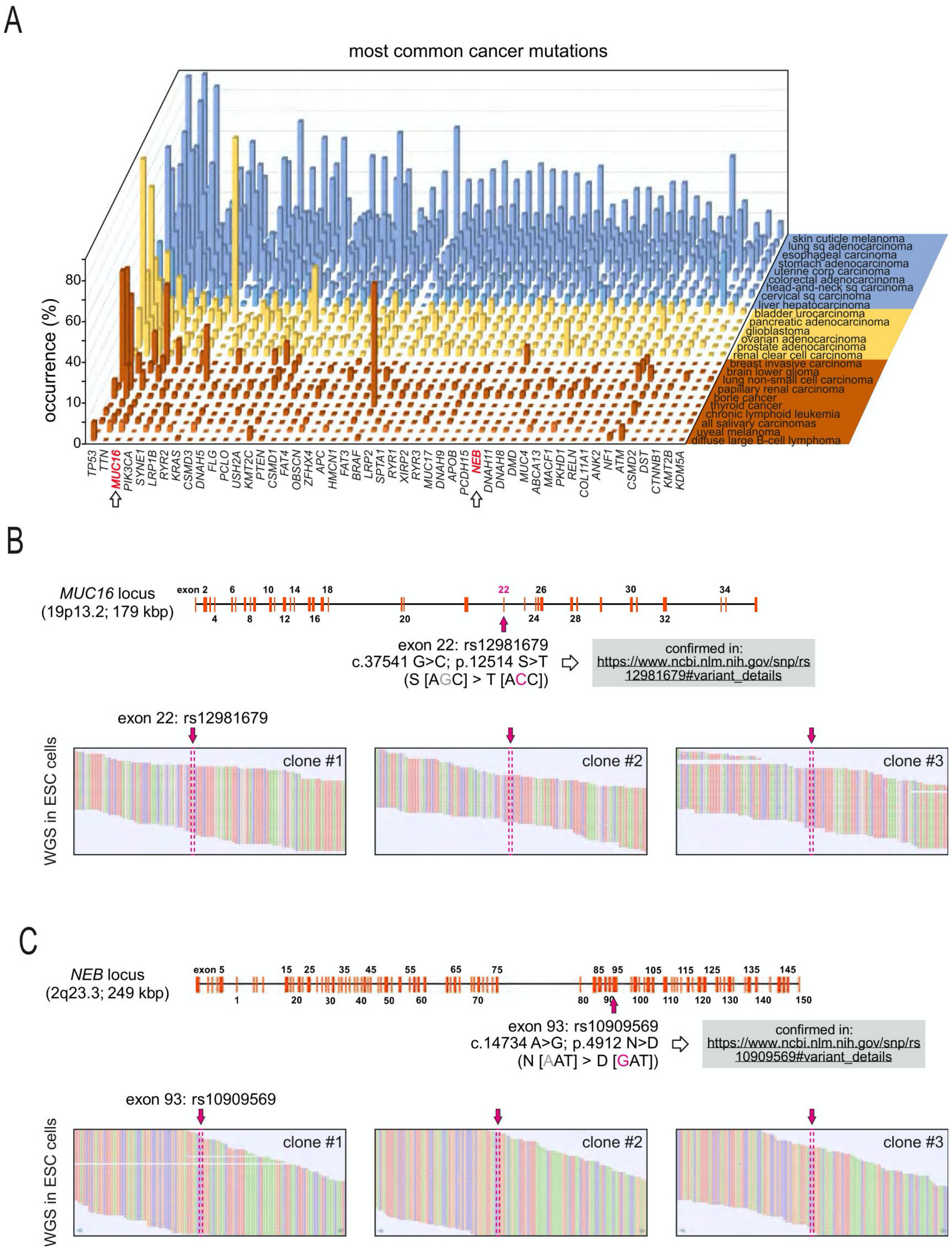


**Figure S4. *CDC6* overexpression recapitulates two of the most frequent cancer gene mutations in ESC cells.**

(**A**) Two of the top 50 most frequent mutations observed in cancer specimens in *MUC16* and *NEB* (*arrows*) were consistently recapitulated in our CDC6-driven cancer evolution model.

(**B**) *MUC16* encodes an established biomarker for diagnosis of many cancers, including lung (the origin of our HBEC model). The identified mutation maps to exon 22 (*arrow*) in a domain associated with protein stabilization and previously confirmed (see SNPdb: rs12981679).

(**C**) As in panel B, but for the *NEB* locus encoding the actin-binding protein nebulin with a mutation in exon 93 also previously confirmed in cancer (see SNPdb: rs10909569).

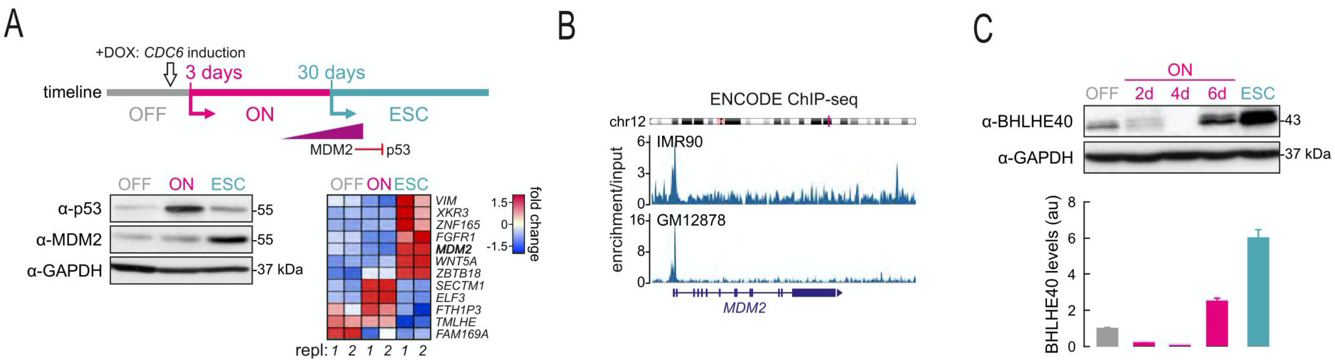


**Figure S5. MDM2 is a BHLHE40 target upregulated in ESC cells.**

(**A**) Western blots and RNA-seq data confirm *MDM2* upregulation and p53 suppression in ESC cells.

(**B**) Genome browser views of BHLHE40 ENCODE ChIP-seq data from IMR90 and GM12878 cells showing strong binding to the TSS of *MDM2*.

(**C**) Western blots showing changing BHLHE40 levels in OFF, ON and ESC cells. The bar graph (*below*; au: arbitrary units) quantifies these changes (mean ±S.D.; n=3).

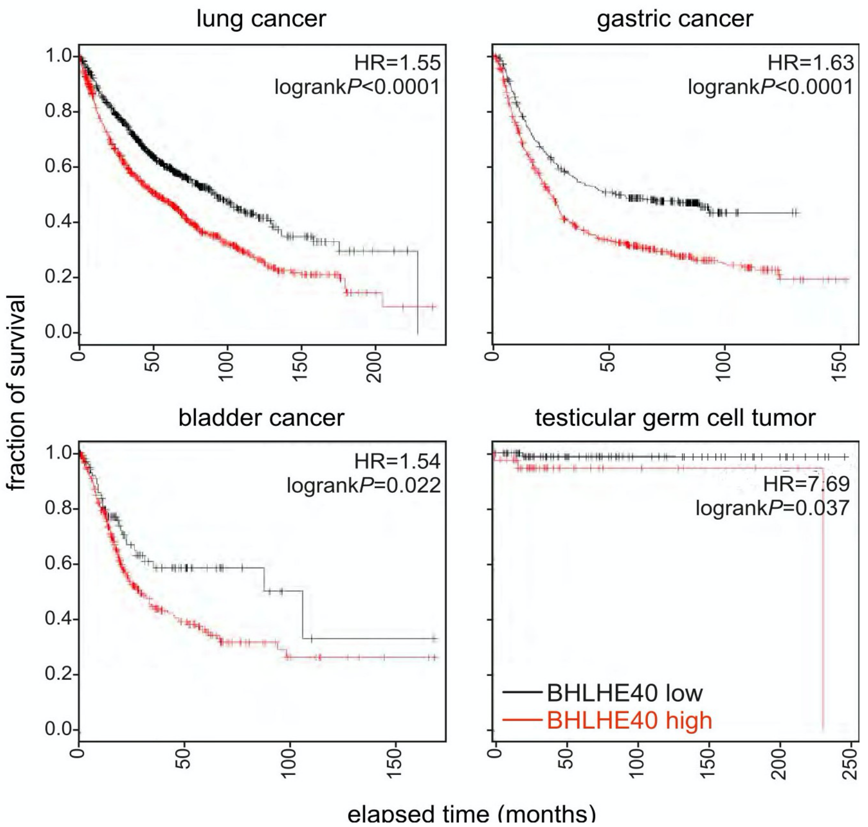


**Figure S6.** ***BHLHE40*** **overexpression in malignancies is associated with poor survival.** Kaplan-Meier survival plots generated using available data (http://www.kmplot.com) from tumors stratified as "high" (red) or "low" CDC6-expressing.

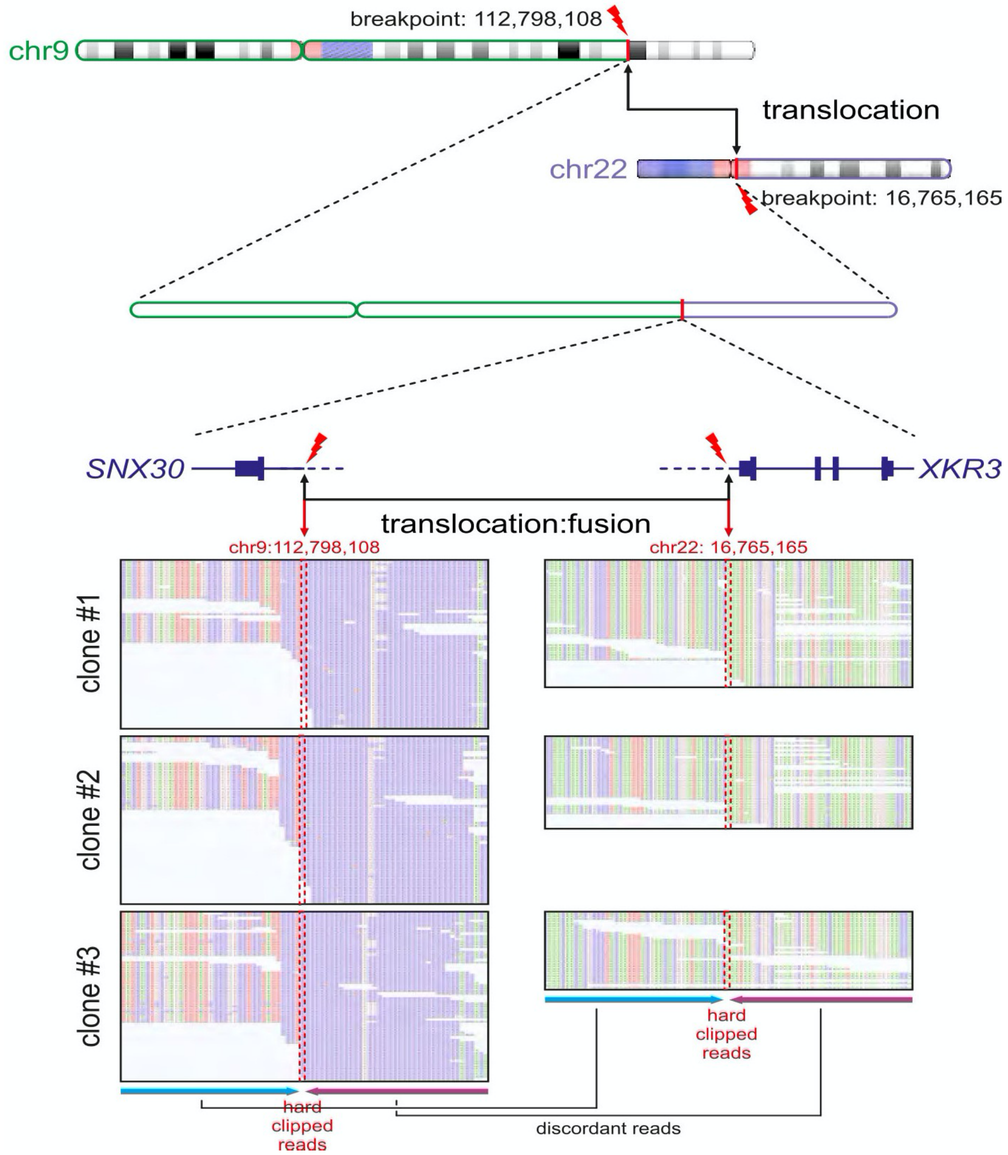


**Figure S7. A reciprocal translocation involving chr9 and 22 among the ESC-shared CNVs.** WGS data describing the translocation breakpoints in ESC cells connecting chr9 and 22. Hard clipped and discordantly mapped reads are indicated for all three replicates.

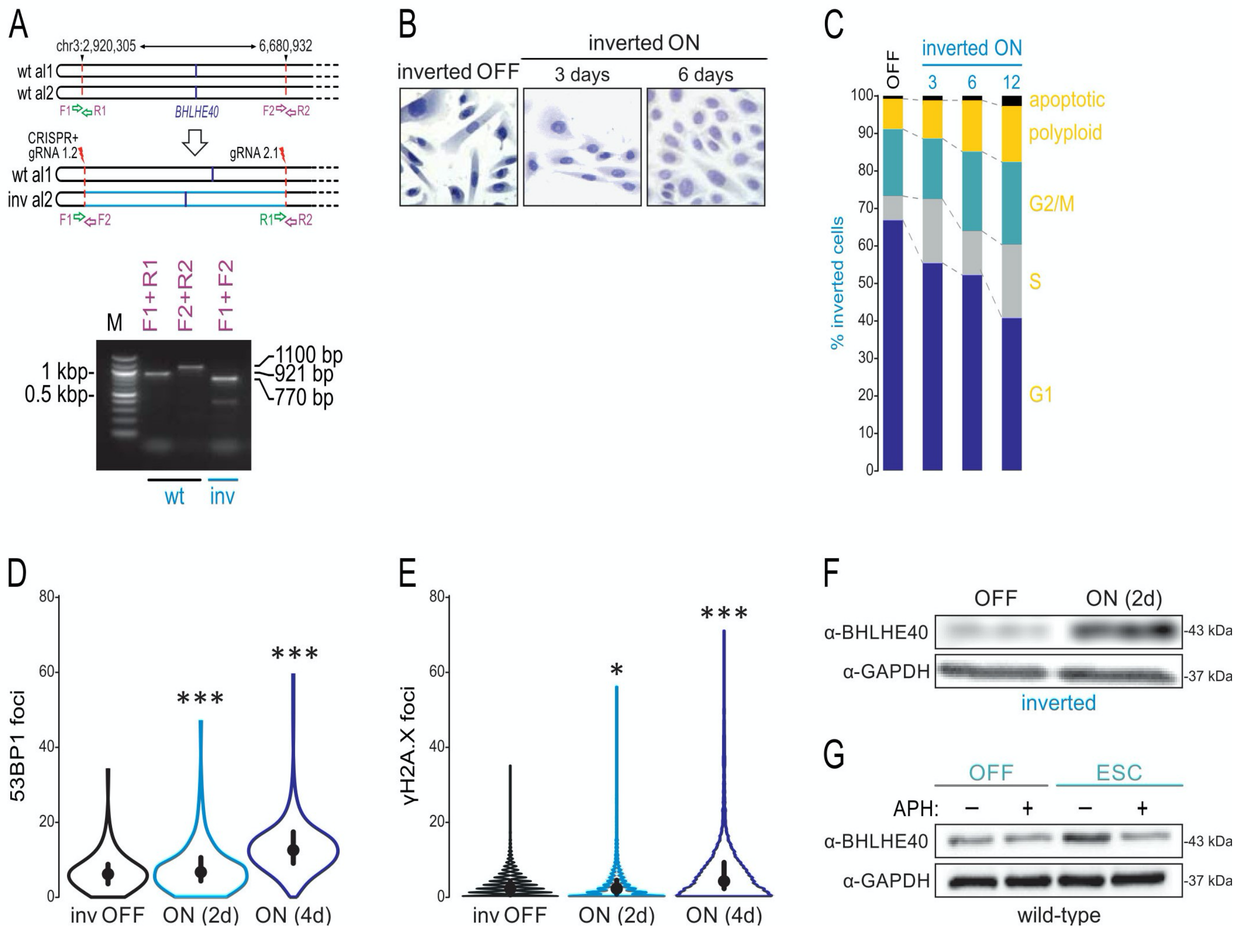


**Figure S8. A second CRISPR-generated clone that harbors the chr3 inversion also bypasses CDC6-induced senescence.**

(**A**) PCR and Sanger sequencing validation of a second clone carrying a CRISPR-generated 3.7-Mbp heterozygous inversion in chr3 that closely mimics that discovered using WGS.

(**B**) Representative images of OFF and 3-/6-day ON "inverted" cells stained with SenTraGor and demonstrating senescence-bypass.

(**C**) FACS analysis of this "inverted" clone indicating increasing S-phase at different days after *CDC6* induction.

(**D**) Violin plots depicting 53BP1 foci accumulation upon *CDC6* induction in "inverted" cells. *: significantly different to OFF; $P < 0.05$, ***: significantly different to OFF; $P < 0.001$, unpaired two-tailed Student's t-test (±S.D.; n=3).

(**E**) As in panel D, but for γH2A.X foci.

(**F**) Western blots showing BHLHE40 overexpression upon *CDC6*-induction; GAPDH provides a loading control.

(**G**) As in panel F, but showing the effect of amphidicolin (APH) treatment on BHLHE40 levels in wild-type OFF and ESC cells.

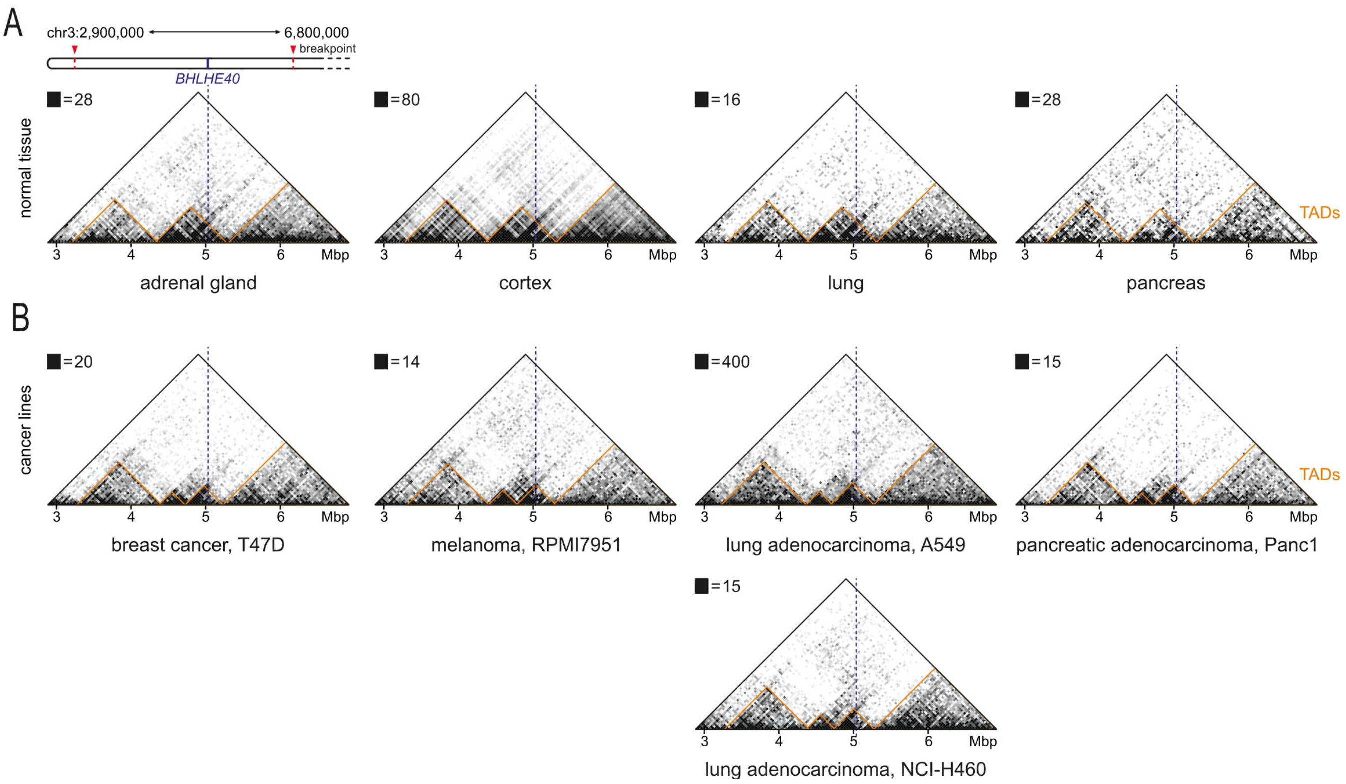


**Figure S9. TAD organization in the extended *BHLHE40* domain.**
(**A**) Hi-C interactions using public data from four normal tissues via the 3D Genome browser (http://promoter.bx.psu.edu/hi-c/view.php) in the 3.9 Mbp around *BHLHE40* (*dotted line*) at 40-kbp resolution. The positions of TADs (*orange triangles*) are also indicated.
(**B**) As in panel A, but for the indicated cancer cell lines.

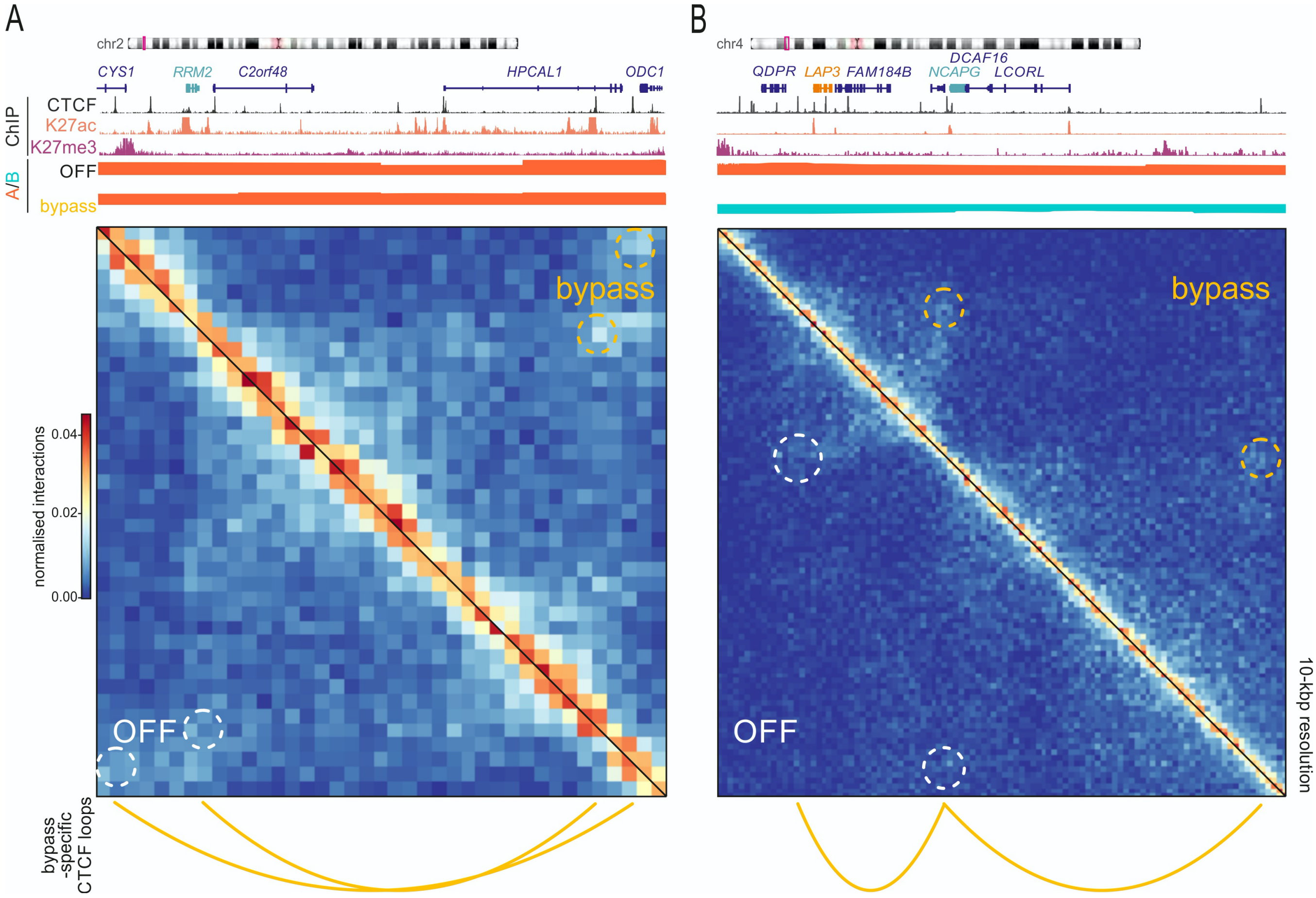


**Figure S10. Changes in 3D chromatin looping explain *CDC6*-induced gene deregulation.**
(**A**) Hi-C heatmaps showing interaction changes between OFF and bypass “inverted” cells in the extended *RRM2* locus on chr2 are shown aligned to CTCF, H3K27ac and H3K27me3 ChIP-seq data, A/B-compartments, and CTCF loop positions. Bypass-specific loops emerging are indicated (*dotted circles*).
(**B**) As in panel A, but for the *LAP3*/*NCAPG* locus on chr4.

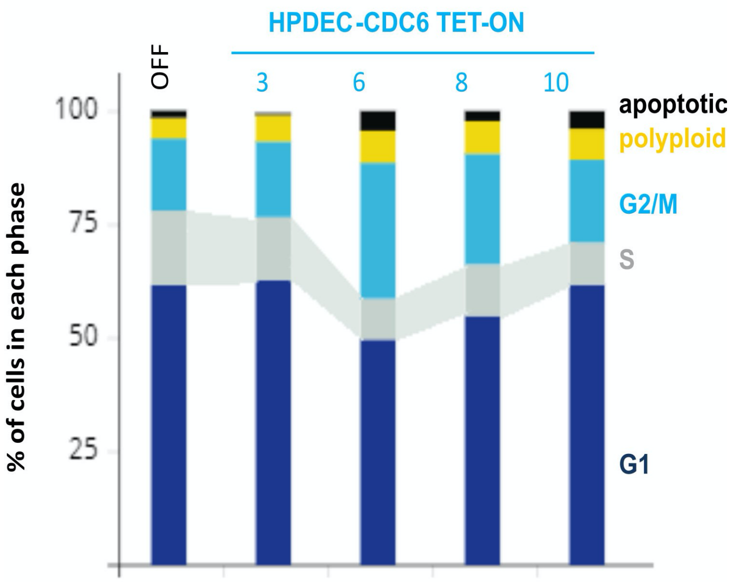


**Figure S11. Cell cycle analysis in an HPDEC-based *CDC6*-TetON system.**
FACS-based cell cycle analysis of HPDECs demonstrating no absence of S-phase at different days after *CDC6* induction.

### SUPPLEMENTAL TABLES AND VIDEOS

**Table S1.** (**A**) Differentially expressed genes located in Common Fragile Sites (CFSs). (**B**) Structural variations (SVs) located in CFSs.

**Table S2.** (**A**) Differentially expressed genes located in copy number variations (CNVs) (CNV ± 20kb). (**B**) Differentially expressed genes within common CNVs and their potential functional implications.

**Table S3.** Differentially expressed genes (DEGs) in "escape" cells that are BHLHE40 transcriptional targets.

**Table S4.** General metrics of Hi-C replicate sequencing and analysis.

**Table S5.** Literature based evidence for *RRM2*, *LAP3*, *NCAPG* overexpression and cancer progression.

**Table S6.** (**A**) DEGs in HBEC *CDC6*-TetON 6-days cells vs bypassed cells that harbor the inversion. (**B**) DEGs in inverted OFF cells versus inverted bypassed cells.

**Table S7.** gRNAs and primer sequences used for the CRISPR/Cas9-mediated generation and validation of the clones harboring the inversion.

**Video S1A-B.** Escape and division (*green circles*) versus senescent/non-dividing (*red circles*) HBEC-CDC6 Tet-ON cells.
